# Epigenetic and brain age across development: Performance and associations in the MIND consortium

**DOI:** 10.64898/2026.07.24.739162

**Authors:** Marlene Staginnus, Vilte Baltramonaityte, Isabel K. Schuurmans, Sarina Abrishamcar, Martin Bauer, Sintia Belangero, Elisabeth B. Binder, Rodrigo A. Bressan, S. Alexandra Burt, Claudia Buss, Shi Yu Chan, Valentine Chirokoff, Shaunna Clark, H. Valerie Curran, Darina Czamara, Serena Defina, Kirsten Donald, Jules R. Dugré, Sonja Entringer, Johan G. Eriksson, Janine F. Felix, Peter Fransquet, Tom P. Freeman, Rodrigo Grassi-Oliveira, Sorcha Hamilton, Christine Heim, Chanelle J. Hendrikse, Anke Hüls, Luke W. Hyde, Natasha S. Jones, Scott A. Jones, Vera N. Karlbauer, Hasse Karlsson, Linnea Karlsson, Nastassja Koen, Will Lawn, Cleanthis Michael, Colter Mitchell, Christopher S. Monk, Michael A. Mooney, Ryan L. Muetzel, Joel T. Nigg, Daniel A. Notterman, Kieran J. O’Donnell, Yi Ying Ong, Vanessa K. Ota, Pedro M. Pan, Tiina Paunio, Jennifer H. Pfeifer, Hung Pham, Jean-Baptiste Pingault, Elmo P. Pulli, Jerod Rasmussen, Leonardo M. Rothmann, Peter A. Ryabinin, Giovanni Salum, Katherine Sawyer, Tim J. Silk, Amalia M. Skyberg, Jolinda Smith, Dan J. Stein, Ai Peng Tan, Ai Ling Teh, Henning Tiemeier, Christopher D. Townsend, Ryan Tung, Jetro J. Tuulari, Pathik D. Wadhwa, Dennis Wang, Catherine J. Wedderburn, Jo Wrigglesworth, Heather J. Zar, Terry Zhou, Charlotte A. M. Cecil, Esther Walton

## Abstract

Understanding how biological age measures perform across development lays the groundwork for investigations into lifespan trajectories of healthy aging. We provide the most comprehensive assessment of epigenetic and brain age models across development (birth to 24 years; ≤20,917 observations across 15 cohorts), evaluating how these models associate with chronological age and with each other, and how these associations change across development. Chronological age-prediction accuracy of epigenetic and brain age models was modest and varied substantially. Accuracy improved with age and stabilized by middle childhood. Few brain and fewer epigenetic clocks performed stably and well across all developmental stages. Performance was better when age range and tissue corresponded between training and testing data. Associations between epigenetic-brain age residuals were small, and changed little across development, tissues or clock generation. Given this developmentally dynamic system of epigenetic-brain age performances and associations, we give key recommendations to improve developmental research in this field.

## INTRODUCTION

Age is an important predictor for various physical, psychiatric, and neurodegenerative disorders^1^. Crucially, individuals with the same chronological age can differ widely in their health status and risk for age-related conditions^2,3^, highlighting the need for a better understanding of pathways underlying healthy aging. Studies have shown that biologically-defined age measures such as DNA methylation (DNAm)-based clocks (i.e., “methylation age” or “epigenetic age/clocks”) or Magnetic Resonance Imaging (MRI)-based brain age might be more accurate in predicting health outcomes than chronological age itself^4^. Specifically, older epigenetic/brain age relative to chronological age (i.e., advanced epigenetic/brain age) is associated with adverse health outcomes, including psychiatric and neurodegenerative disorders, reduced cognitive functioning, and mortality, whereas a younger biological age may provide insights into delayed developmental processes and resilience to age-related decline^5–8^.

Despite an increasing body of research on aging and aging-related biomarkers, previous studies have predominantly trained and applied these biomarkers in adult populations. However, (biological) aging starts in utero and many determinants of healthy aging are evident in early life^9–11^. Although researchers are beginning to apply these biological age measures to pediatric populations, it is unclear to what degree these models are valid in - and portable to - earlier developmental stages^12–14^, hindering efforts to understand how healthy aging trajectories emerge across development. While chronological age prediction is not the primary objective of all biological age models – particularly next-generation clocks (Table 1) – correspondence with chronological age functions as a key benchmark for comparing model performance across developmental stages before assessing associations with health-relevant traits.

**Table 1.**
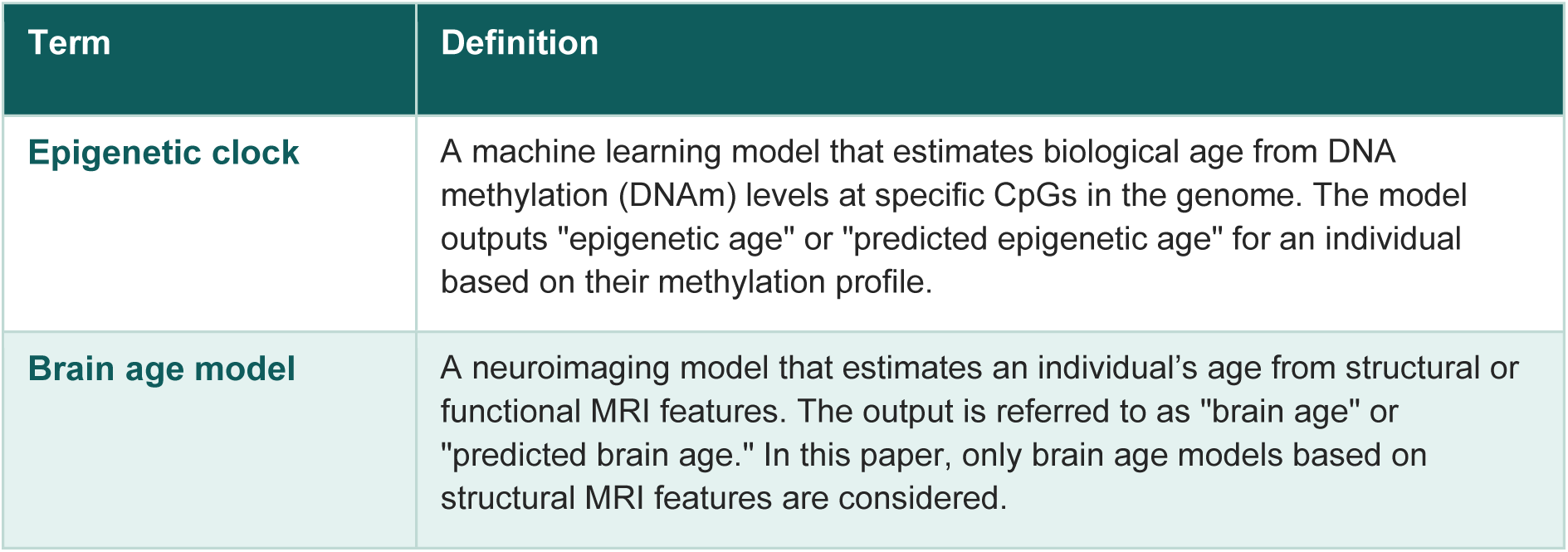
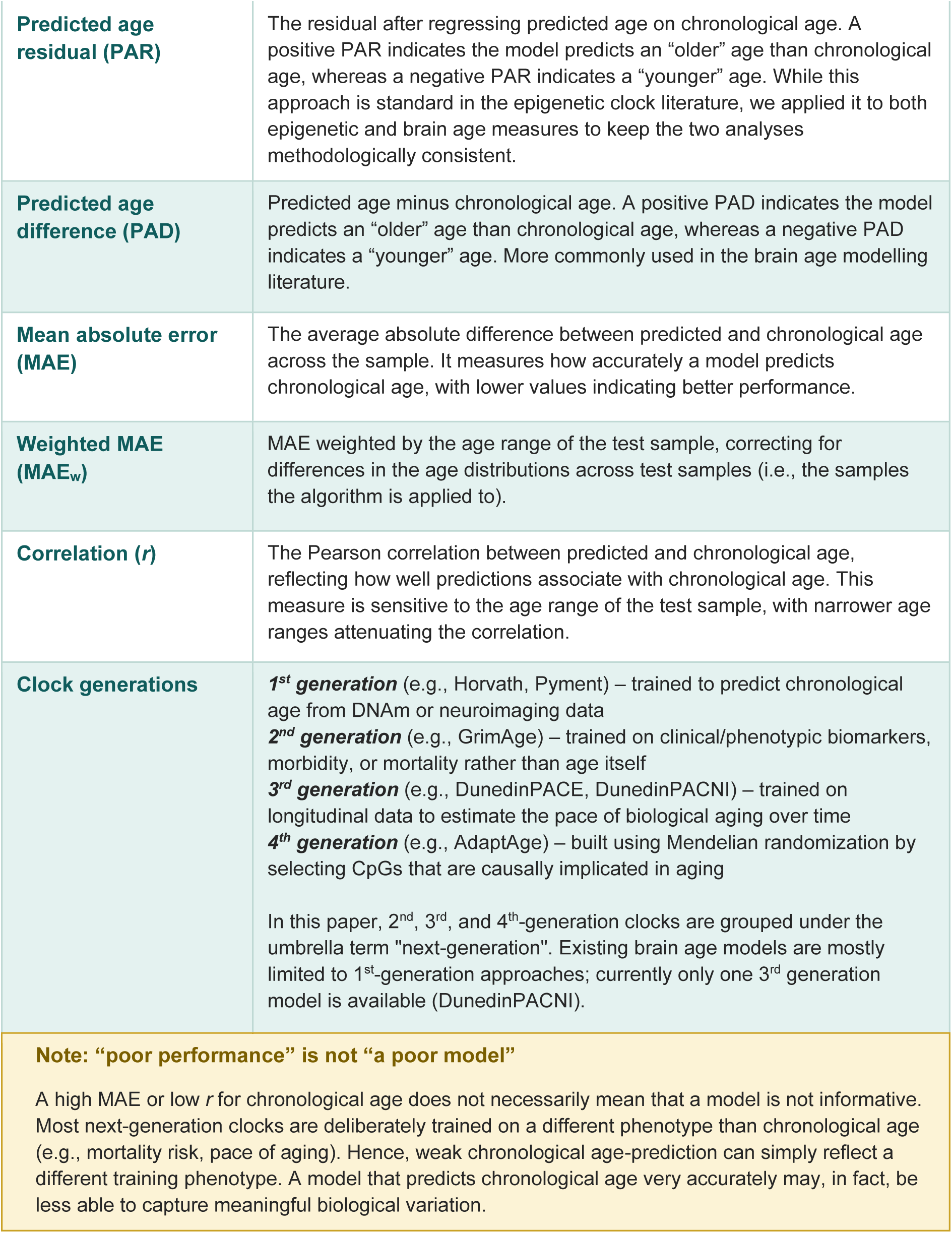
Glossary of key terms.

It furthermore remains unclear how different epigenetic and brain age measures relate *to each other* across development. Aging has been defined as a system-wide process^15,16^, suggesting potential intercorrelations between different epigenetic and brain age measures. Conversely, studies have observed substantial variability in aging rates across different tissue types^17,18^, denoting potentially distinct epigenetic and brain aging trajectories that might differ in how they predict health outcomes later in life. As many brain-associated disorders have a developmental origin^19^, knowing how epigenetic and brain age measures relate to each other in development is needed to inform risk prediction and early intervention efforts.

Of the few studies investigating epigenetic and brain age measures jointly, three gaps remain. First, these studies have primarily investigated epigenetic age and brain age as separate predictors of health outcomes, rather than examining the extent to which they relate to each other (e.g.,^20^). Second, most studies correlated epigenetic age measures with only a *single* brain age model (e.g.,^21–23^) despite key differences between models (e.g., different training sample characteristics or imaging features). Third, and most importantly, these studies did not investigate child and adolescent samples, even though epigenetic patterns and brain structure change at different rates across development^24–26^, highlighting the need to examine the association between epigenetic and brain age measures across several developmental stages.

To address these gaps, we provide the most comprehensive assessment of how a wide range of epigenetic and brain age models relate to chronological age from birth to young adulthood and how they associate with each other. We also explore what features (e.g., developmental period, tissue) might explain heterogeneity in performance and associations; whether certain epigenetic clocks more strongly associate with brain age than others; and how these cross-sectional associations change across development.

As preregistered, we hypothesized that epigenetic and brain age models would perform better in samples where the age range overlaps with that of the model’s training sample. Furthermore, we hypothesized that first-generation epigenetic clocks would show a stronger association with first-generation brain ages compared to next-generation epigenetic clocks as the former are both trained to predict chronological age. Third, we predicted that epigenetic age measures trained in brain tissue – or tissues such as saliva that show a DNAm profile more similar to brain than blood ^27^ – would associate more strongly with brain age measures. We did not make predictions on how these associations might differ across development.

Accompanying this study, we also provide a fully searchable dashboard (https://epi-brain-age-dashboard.streamlit.app/) to support researchers in identifying the most suitable biological age measure(s) for their developmentally-focused studies.

## METHODS

### Cohorts

This preregistered study (https://doi.org/10.17605/OSF.IO/6CSM3) drew on cohorts from the Methylation, Imaging, and NeuroDevelopment (MIND) Consortium^28^, which aims to strengthen the developmental focus in the field of neuroimaging epigenetics. The consortium consists of cohorts from Europe, North and South America, Africa, Australia, and Asia, which have DNAm and MRI data for at least one (but mostly multiple) timepoints between birth and young adulthood (Figure 1; SM Table S1). For further details (such as inclusion/exclusion criteria) and for deviations from our preregistration, see SM 1.1 and 1.2.

**Figure 1.**
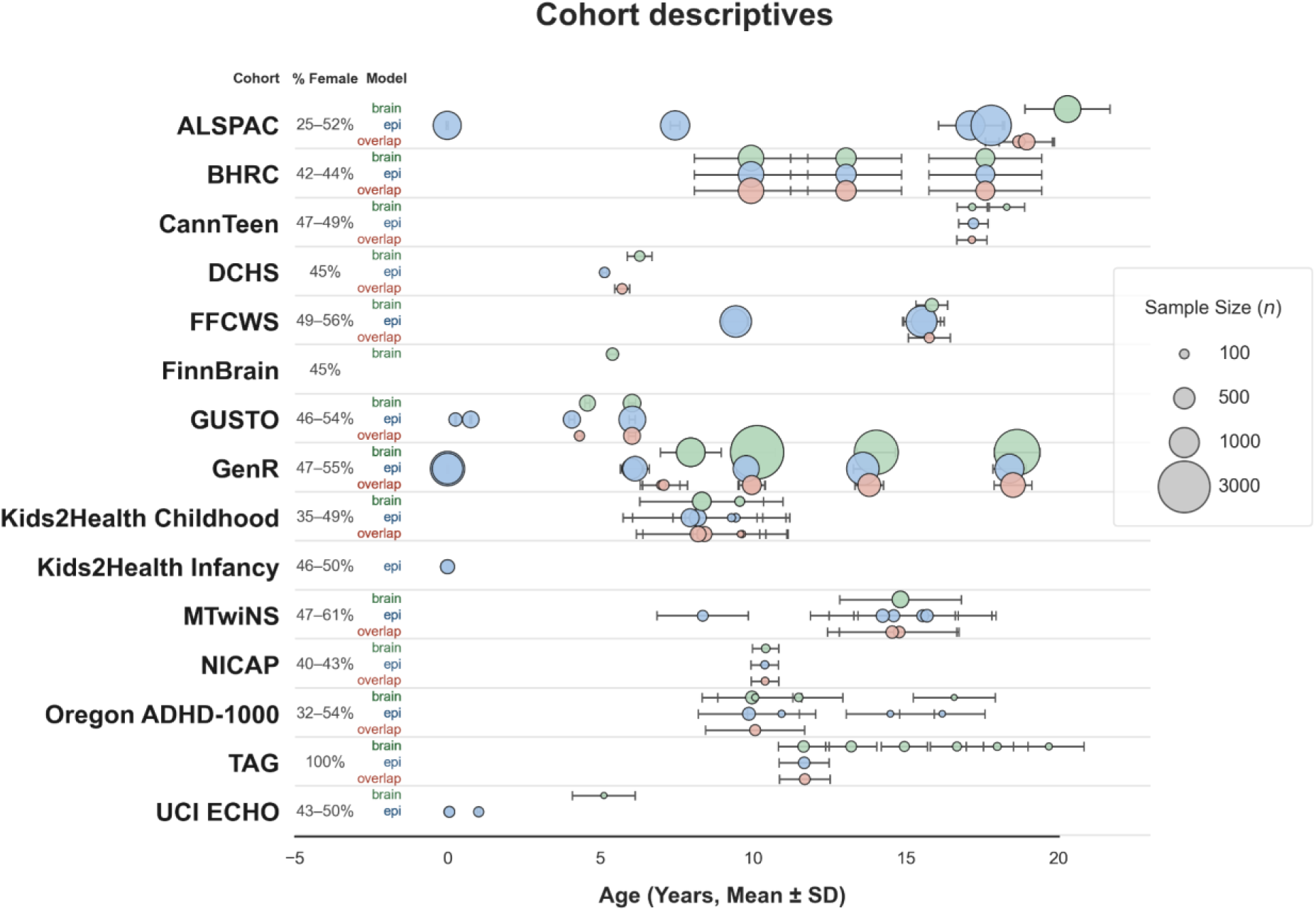
Cohort descriptives. Number of timepoints per developmental age bracket (if a cohort has two different tissues or arrays at a specific timepoint, this was counted as one timepoint): Epigenetic: birth = 4, infancy = 3, early childhood = 2, middle childhood = 10, late childhood = 7, adolescence = 6, young adulthood = 1; brain: early childhood = 3, middle childhood = 7, late childhood = 10, adolescence = 6, young adulthood = 4, overlap (timepoints with both brain and epigenetic data available): early childhood = 2, middle childhood = 7, late childhood = 5, adolescence = 4, young adulthood = 1.

### Measures

#### DNA methylation

##### DNAm preprocessing

DNAm levels obtained from blood (cord, whole), placental, buccal or saliva tissues were assessed using the Illumina Infinium HumanMethylation450K BeadChip assay (Illumina 450 K array) and Infinium MethylationEPIC (v1 and v2) BeadChip (Illumina, San Diego, CA, USA). For detailed information on DNAm preprocessing, see https://github.com/MarleneSt/MIND_BrainAge-EpiAge, SM 1.3 and SM Table S3.

### Epigenetic age models

To ensure comprehensiveness, we carried out a scoping search for publicly available epigenetic age models and assessed their feasibility. Models were included if they were publicly available and if the age range of the sample used for model training overlapped at least partly with the age range investigated in this study (0-24 years). We also included DunedinPACE even though DNAm training data was collected from adults aged 45 years. This was due to the uniqueness of its trained phenotype (i.e., longitudinal rates of biological aging measured from age 26 to 45 years) and outcome measure (pace of aging). In total, we included 20 clocks (SM Table S2), covering 14 first-generation clocks (including three gestational age clocks), three second-generation clocks, one third-generation clock and two fourth-generation clocks (Table1).

### Neuroimaging

#### MRI preprocessing

Structural T1-weighted MRI scans were acquired using 1.5 or 3T scanners and preprocessed using FreeSurfer^29^ version 6 or higher. For further details, see https://github.com/MarleneSt/MIND_BrainAge-EpiAge, SM 1.4 and SM Table S4.

#### Brain age models

Our scoping review identified 15 potential brain age models. Of those, we included models that i) were publicly available, ii) had a training sample age range that at least partly overlapped with the age range investigated in this study, iii) focused on structural T1-weighted MRI scans, and iv) relied on FreeSurfer or minimal preprocessing (to ensure feasibility). Consistent with our rationale to include the epigenetic age clock DunedinPACE, we also included DunedinPACNI here despite a training sample age of 45 years. In total, we included eight models (SM Table S2), covering seven first-generation models and one third-generation model (Table 1).

### Analyses

Cohort-level analyses were performed cross-sectionally per timepoint covering birth to young adulthood, and then meta-analyzed across timepoints, using inverse-variance weighted multi-level random-effects meta-analyses using the *metafor* R package^30^. Models accounted for nesting in cohorts, timepoints, and individual age model (or brain age-epigenetic age model pairs, respectively).

To answer how well epigenetic clocks and brain age models perform in development, we focused on i) the Mean Absolute Error (in relation to chronological age), weighted by the test sample age range (MAE_w_) to increase comparability across samples; and ii) Pearson’s correlation coefficient with chronological age. Moreover, we considered eight additional metrics including unweighted MAEs (SM 1.5 and https://epi-brain-age-dashboard.streamlit.app/). To address how brain and epigenetic age measures associate with each other, we performed robust linear regression analyses using an M estimator (with heteroscedasticity consistent standard errors) with the brain-predicted age residual (brain-PAR, residualized against chronological age) as the outcome and epigenetic-predicted age residual (epigenetic-PAR) as the predictor. PAR was chosen for both epigenetic and brain age as this measure is age-independent and allows for better comparisons across cohorts with different age structures^31^.

Chronological age was calculated in years (≥2 decimals). To facilitate integration of gestational clocks (which produce predicted age in gestational weeks) into meta-analyses, these were also transformed into years (SM 1.6).

Our primary model corrected for age at MRI, age at DNAm (if correlated < 0.8 with age at MRI), sex, and batch. Additional models were run 1) without covariate adjustment, 2) primary model covariates and estimated cell type proportions (SM 2.12), and 3) when restricting to model combinations with 100% training and test sample age range overlap and where epigenetic clocks were applied to their intended tissue (SM 2.13). Moderator analyses explored the effects of model or model pair (i.e. obtaining specific results for individual clocks and model association pairs), age, proportional overlap of training and test age range, tissue (epigenetic only), processing level (MRI only), and epigenetic clock generation. Results were false discovery rate (FDR)-corrected (across moderator analyses per outcome and across all models/model-combinations, respectively) and leave-one-cohort-out analyses were performed (reported throughout the SMs). For further details on our statistical approach, see SM 1.7.

Results can also be queried through our dashboard (https://epi-brain-age-dashboard.streamlit.app/).

## RESULTS

### Cohort descriptives

A total of 15 MIND cohorts contributed data from birth to mean age 20 years (age_max_ = 24; Figure 1). Twelve cohorts had repeated measures available at up to six points, while three cohorts contributed data obtained at a single timepoint. Epigenetic data were available for 20,917 observations (including repeated measures) across 33 timepoints (birth to 18 years) while MRI data were available for 19,925 observations across 30 timepoints (4 to 20 years).

Overlapping data were available in 5,559 observations across 19 timepoints (4 to 20 [MRI] and 18 [epigenetic] years, SM Table S5-S8). For a breakdown per developmental age bracket, see Figure 1 note. All but one cohort (TAG, 100% female) were sex-mixed with the proportion of female participants ranging between 24.6-60.6%.

### Performance of epigenetic clocks across development

The unweighted pooled MAE was 6.61 years (95% CI [2.78–15.70], SM 2.1). The overall pooled MAE_w_ (weighted by the test sample age range) across all epigenetic clocks, cohorts and developmental periods was 2.18 (95% CI [0.74, 6.43], Figure 2A, SM Table S9), indicating that the error was on average 2.18 times larger than the sample age range. In line with this wide performance range, we observed very high levels of heterogeneity (I^2^ = 99.95%, *p* < .001), that was mostly attributable to clock models (60.68%), followed by cohorts (29.84%) and timepoints (within cohorts; 9.44%). Performance differed significantly between models (QM(18) = 225166.02, *p*_FDR_ < .001) with MAEs_w_ ranging between 0.03 (Bohlin) and 27.34 (DamAge, Figure 2A). Fourth-generation clocks showed the highest MAE_w_, followed by second-generation clocks, while first-generation clocks (especially gestational clocks) showed the lowest MAE_w_ (SM 2.1) consistent with first generation clocks being trained to predict chronological age and gestational clocks only being applied at birth timepoints. These patterns remained similar when considering correlations between chronological and predicted age. Correlations were overall moderate in magnitude (*r* = .35, 95% CI [.17, .50], SM 2.1, Figure 2B, SM Table S10), likely a consequence of the narrow sample age ranges.

**Figure 2.**
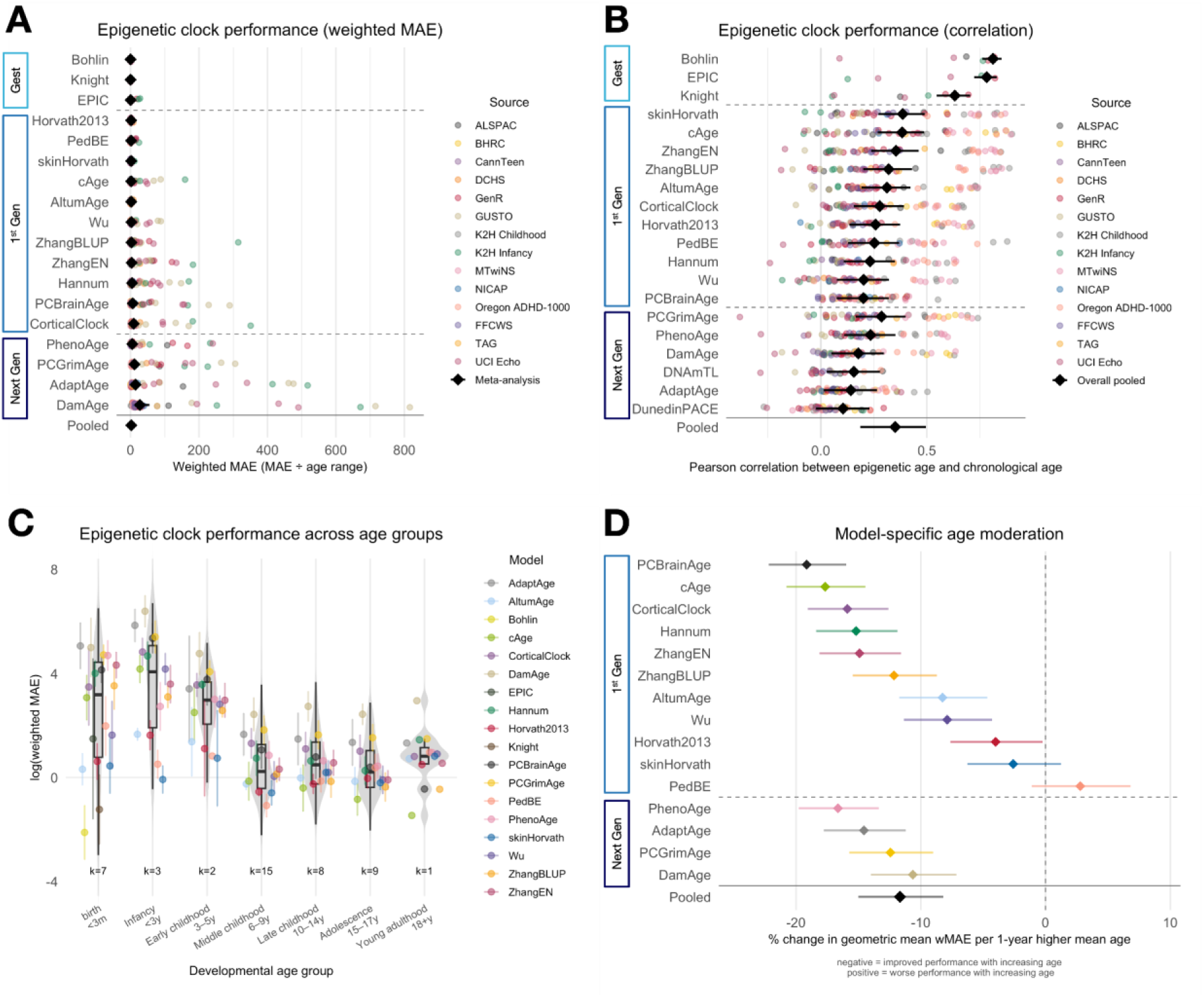
Epigenetic clock performance. A) Epigenetic clock performance based on the mean absolute error weighted by the test sample age range (MAE_w_) ordered by clock generation and performance (best to worst). Two epigenetic clocks (DNAmTL and DunedinPACE) were not included in MAE_w_ analyses as this performance measure is not meaningful for these models as neither of them produce predicted age in years. B) Clock performance based on the correlation with chronological age (Pearson’s correlation coefficient) ordered by clock generation and performance (best to worst). Correlations between chronological age and DNAmTL predictions were inverted prior to meta-analyses as it predicts telomere length, where greater values = younger. C) Performance based on log(MAE_w_) across age groups. Meta-analyses per age group were performed for visualization purposes only, whereas the age-related analyses reported in the main text were continuous. D) Performance stability over development (0: no change with age, >0: poorer performance with age, <0: better performance with age). Gest=1st generation clocks trained to predict gestational age. Gen=Clock generation whereby second to fourth generation clocks are considered ‘Next Gen’. k= the number of unique cohort–timepoint combinations within an age bin (if a cohort has two different tissues or arrays at a specific timepoint this will be counted as two timepoints). These plots can be explored in greater detail in our dashboard: https://epi-brain-age-dashboard.streamlit.app/.

### Factors that influence epigenetic clock performance

#### Developmental period

Epigenetic clock age-prediction accuracy (excluding gestational clocks) improved with increasing age (*b_l_*_og_ = -0.12, *p*_FDR_ < .001, yearly decrease in MAE_w_ of 11.67%) and stabilized by middle childhood (Figures 2C and 2D). The magnitude of these improvements differed modestly between clock generations, with second-generation clocks showing the steepest age-related reductions in prediction error (SM 2.2). Most epigenetic clocks (e.g., PCBrainAge, cAge, PhenoAge and CorticalClock) showed substantially better prediction performance with age, but some clocks (e.g., PedBE, Horvath2013, skinHorvath) were comparatively stable across development (Figure 2D, SM Table S9).

In contrast to MAE_w_, the overall correlation between epigenetic-predicted age and chronological age did not significantly change with age (*b*_z_ = 0.01, *p*_FDR_ = .154), although as before, we observed an increase in correlation by middle childhood and age-dependent correlation changes in individual clocks (SM 2.2, SM Table S10). DunedinPACE (*b*_z_ = -0.01, *p*_FDR_ = .029) showed a significant negative age slope, reflecting decreasing correlations with chronological age in older samples.

#### Age range overlap between training and testing data

As hypothesized, we found that a greater proportional overlap between the training and testing age range was associated with significantly lower MAE_w_ (*b*_log_ = -0.01, *p_FDR_* < .001). The predicted geometric mean MAE_w_ decreased from approximately 5.95 at 0% overlap to 3.04 at 100% overlap, indicating progressively better apparent epigenetic age-prediction accuracy with increasing age overlap. Age-correlations showed the same effect (*r*_no.overlap_ = .18 to *r*_overlap_ = .35, *b*_z_ = 0.002, *p_FDR_* < .001; SM 2.3).

#### Tissue match between training and testing data

Epigenetic age models performed significantly better when the tissue type of the test sample matched the tissue(s) used during model training (*b*_log_ = -0.22, *p_FDR_* < .001). Predicted geometric mean MAE_w_ was 2.41 in non-matching samples versus 1.92 in matching samples. Age-correlations increased from *r* = .31 to *r* = .42 between non- to matching tissues (*b*_z_ = 0.11, *p_FDR_* < .001; SM 2.4).

### Performance of brain age models across development

The unweighted MAE across all brain age models was 4.46 years (95% CI [2.48, 6.43], SM 2.5). The overall pooled MAE_w_ was 1.72 (95% CI [0.67, 2.77]), indicating an error 1.72 times larger than the sample age range (Figure 3A). There was high heterogeneity (I^2^ = 99.98%, *p* < .001), but this time it was mainly accounted for by cohort (94.40%), while only some (5.42%) or almost no variation (0.19%) were attributable to model and developmental timepoints, respectively. Performance differed significantly between models (QM(7) = 7699.08, *p*_FDR_ < .001; Figure 3A). Pyment and DevBrainAge showed the best MAEs_w_ (SM Table S11). Similarly, Pyment showed the strongest correlation with age (*r* = .34, 95% CI [.25, .40]), while DunedinPACNI was the only model showing a significant negative correlation (*r* = -.10, 95% CI [-.19, -.02], Figure 3B, SM Table S12). The overall age-correlation across all brain models, cohorts and developmental periods was small-to-moderate in magnitude (*r* = .20, 95% CI [.07, .32], SM 2.5), likely a reflection of the narrow age range in many samples.

**Figure 3.**
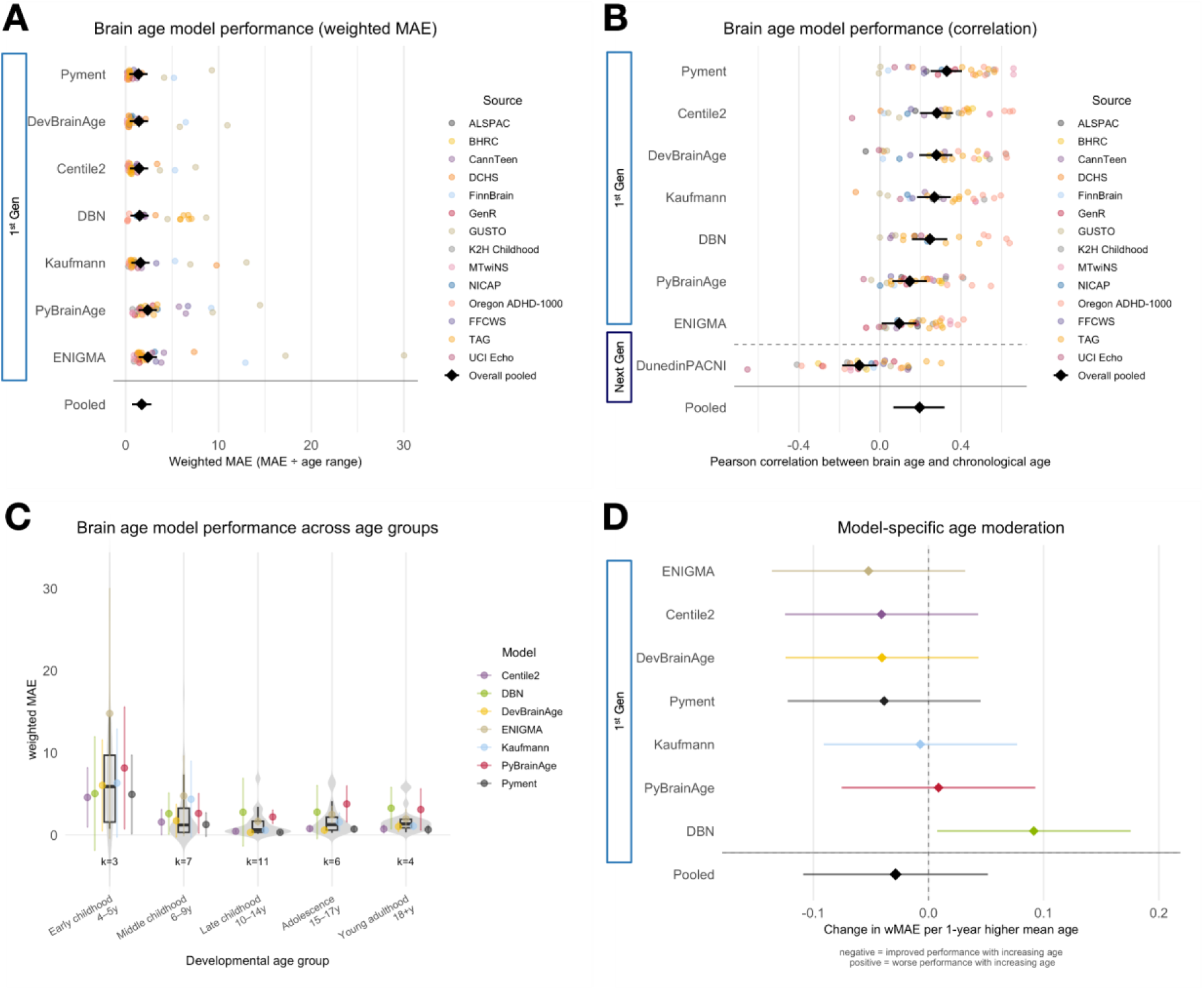
Brain age model performance. A) Brain age model performance based on the mean absolute error weighted by the test sample age range (MAE_w_) ordered by performance (best to worst). One brain age model (DunedinPACNI) was not included in MAE analyses as this performance measure is not meaningful here as the model does not produce predicted age in years. B) Brain age model performance based on the correlation with chronological age (Pearson’s correlation coefficient) ordered by model generation and performance (best to worst). C) Performance across age groups. Meta-analyses per age group were performed for visualization purposes only, whereas the age-related analyses reported in the main text were continuous. D) Performance stability over development (0: no change with age, >0: poorer performance with age, <0: better performance with age). Gen=Model generation. k= the number of unique cohort–timepoint combinations within an age bin. These plots can be explored in greater detail in our dashboard: https://epi-brain-age-dashboard.streamlit.app/.

### Factors that influence brain age model performance

#### Developmental period

While the overall age effect was non-significant (MAE_w_: *b* = -0.03, 95% CI [-0.11, 0.05], *p*_FDR_ = .484; correlation: *b*_z_ = 0.01, *p*_FDR_ = .242, SM 2.6), we observed a similar pattern as for the epigenetic clocks, whereby performance was slightly better and stabilized by middle childhood (Figure 3C). Additionally, we observed only weak age-related performance changes in individual clocks (Figure 3D, SM 2.6 and SM Table S11+12).

#### Age range overlap between training and testing data

As hypothesized, greater proportional overlap between the cohort age range and the brain model training age range was associated with significantly lower MAE_w_ (*b* = -0.003, *p*_FDR_ < .001). Predicted MAE_w_ decreased from approximately 2.01 at 0% overlap to 1.67 at 100% overlap, indicating progressively better apparent model performance with increasing age overlap (SM 2.7). Correlational results aligned with these patterns (*r*_no.overlap_ = .10 to *r*_overlap_ = .24, *b*_z_ = 0.001, *p*_FDR_ < .001; SM 2.7). For MRI preprocessing effects, see SM 2.8.

### Epigenetic PAR - brain PAR associations

Averaged across all model combinations, we found a small positive association between brain PARs and epigenetic PARs (β = 0.017, 95% CI [0.01, 0.03], *p* = 0.001, Figure 4A), which was much smaller than within-modality associations (covariate-unadjusted *r*_brain.PAR_ = .25, *r*_epi.PAR_ = .22, SM 2.9). While we found no strong evidence for variability in effect magnitude across model combinations, cohorts or timepoints (I^2^ = 14.64%, *p* = 0.578), associations differed significantly between model combinations (QM(136) = 256.92, *p*_FDR_ < .001). Among the 136 model combinations, we observed both negative (22.1%) and positive (77.9%) pooled associations (Figure 4B, SM Table S13). The largest pooled PAR associations were observed between Kaufmann and skinHorvath (β = 0.06, 95% CI [0.03, 0.10]), DevBrainAge and cAge (β = 0.06, 95% CI [0.03,0.10]), and Centile2 and DNAmTL (β = -0.06, 95% CI [-0.121,0.001]).

**Figure 4.**
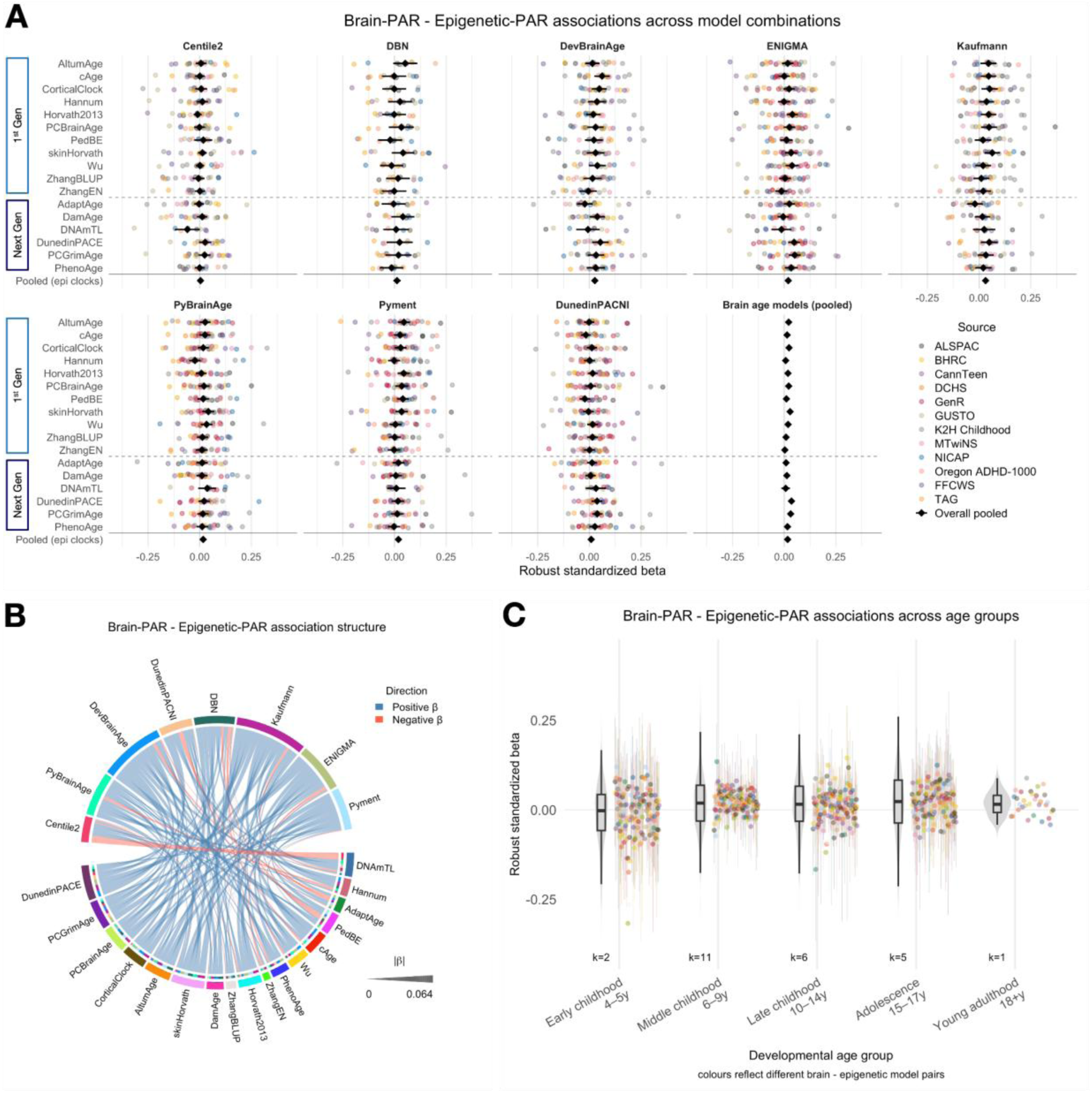
Associations between brain age residuals and epigenetic age residuals. A) Pairwise epigenetic-brain PAR associations. B) Cord diagram showing the pooled strength of associations between all model combinations. C) Associations across age groups with colors reflecting the different model-combinations. Meta-analyses per age group were performed for visualization purposes only, whereas the age-related analyses reported in the main text were continuous. Associations with the epigenetic clock DNAmTL were inverted prior to meta-analyses as it predicts telomere length where greater values = younger. PAR=predicted age residual; β=robust standardized beta; k= the number of unique cohort–timepoint combinations within an age bin (if a cohort has two different tissues or arrays at a specific timepoint this will be counted as two timepoints). These plots can be explored in greater detail in our dashboard: https://epi-brain-age-dashboard.streamlit.app/.

### Factors that influence epigenetic PAR - brain PAR associations

#### Developmental period

The strength of association between epigenetic PAR and brain PAR was not significantly moderated by sample mean age, although the estimated effect was positive indicating that associations became stronger with age (*b* = 0.001, 95% CI [−0.001, 0.003], *p*_FDR_ = .477; Figure 4C).

#### Epigenetic clock generation

Contrary to our hypothesis, epigenetic PAR–brain PAR associations did not significantly differ between first- and next-generation clocks overall (*b* = 0.001, *p*_FDR_ = .700). In exploratory analyses, we also modelled each generation separately. Here, results showed that associations differed across epigenetic clock generations (QM(4) = 20.11, *p*_FDR_ = .002), with the strongest associations observed for DunedinPACE, the only third generation clock (β = 0.033 95% CI [0.017, 0.049], SM 2.10).

#### Tissue

Also contrary to our hypothesis, tissue of the epigenetic clock training and/or testing data did not significantly moderate epigenetic PAR–brain PAR associations (saliva: *b* = 0.004, *p*_FDR_ = .477; buccal: *b* = 0.004, *p*_FDR_ = .477; brain: *b* = 0.007, *p*_FDR_ = .128, SM 2.11).

## DISCUSSION

Chronological age-prediction accuracy of epigenetic and brain age models was often modest, with pooled prediction errors exceeding the age range. Performance differed between models and was better in later developmental stages (especially for epigenetic age) and when tissue or age range overlapped between training and testing data. This indicates that some models are better suited for developmental applications than others when using chronological age as a benchmark. Associations between epigenetic age and brain age residuals were small yet largely positive, remaining stable over time and tissues.

### Epigenetic and brain age performances are highly heterogenous

Although a large degree of heterogeneity and lower performance was expected, especially for next-generation models not trained to predict chronological age, we also found heterogeneity within (non-gestational) first-generation epigenetic clocks and – to a lesser extent – brain age models.

*Recommendation #1: Researchers should preselect clocks that align with their study rationale and cohort characteristics*.

For developmental studies aiming to closely track epigenetic or brain age deviations from chronological age, first-generation clocks might be preferred. For narrow-timepoint data, researchers should i) select clocks that perform well in that developmental period (e.g., Bohlin/Knight at birth, skinHorvath or PedBE in childhood, cAge in adolescence) and ii) prioritize clocks that were trained across a similar age range and tissue as their data. For data covering a wider age range, researchers should select clocks that perform stably over this period (e.g., PedBE or (skin/)Horvath perform well from infancy to young adulthood). For brain age, we recommend the use of models such as Pyment and Centile2 that perform consistently well across developmental stages.

We emphasize that tracking deviations from chronological age (i.e. the focus of this paper) might not always be the key objective. Tracking healthspan, mortality risk, the rate of aging or diving deeper into causal considerations can be alternative aims, where other, next-generation models might perform better than the ones we highlighted above (see also^32–34)^.

### By middle childhood epigenetic (and brain age) models perform better

We found that age-prediction accuracy improved for epigenetic and brain age models stabilizing from middle childhood onwards. This age effect was more pronounced (and only significant) for epigenetic clocks.

*Recommendation #2: Researchers should be cautious in the interpretation of null findings at early developmental stages*.

Age-dependent improvements in prediction could reflect either an emerging association (e.g., in late versus early childhood) that is due to true developmental effects or an artifact caused by poor model performance at the earlier timepoint. In these cases, we recommend using developmentally stable clocks (e.g., PedBE, Pyment). Going forward, new clocks built to explicitly capture these changing developmental patterns (e.g., trained on repeated measures) might prove useful.

*Recommendation #3: We need clocks that are designed for early developmental datasets*.

Although most brain age and some epigenetic models (e.g., Horvath) included developmental timepoints in their training, this was often only a small fraction of the overall data. One of the exceptions is PedBE that was trained exclusively on pediatric samples between age 0 to 20 years, while DevBrainAge was trained in data ranging from 9 to 19 years. Our finding that both models are among the better and more stably performing clocks in development highlights how a developmental focus can improve prediction^14^, but several challenges remain. First, we were unable to apply current brain age models to neuroimaging data in infants due to methodological difficulties (e.g., reversal of light-dark contrast in T1-weighted images, extracting specific imaging features from infant scans). Second, there are currently no next-generation clocks that are built using developmental training data. Progress on these issues is key to understanding the importance of early development on lifelong (brain) health, mortality, and rates of aging. Relatedly, in development both advanced and delayed aging might be associated with adverse effects, which might not be captured by existing clocks that were predominantly trained on adult data.

### Small positive brain PAR-epigenetic PAR associations that change only weakly across developmental stages or tissues

Small associations between brain PARs and epigenetic PARs (even within first-generation models) across development align with previous research that reported a mosaic of aging modalities^35^ and suggest weakly related epigenetic and brain age profiles that might also differ in how they predict health outcomes later in life^36^. Systems might become more interconnected with age (as environmental exposures accumulate), in line with previous research showing epigenetic-brain age links by middle adulthood^37,38^. Weak tissue effects are surprising and suggest that training epigenetic clocks in cortical tissue might not get closer to capturing brain features that constitute brain age. However, the lack of tissue effects could also reflect the overall weak brain PAR-epigenetic PAR association.

*Recommendation #4: Researchers interested in the interconnections between epigenetic and brain aging systems should anticipate that these systems may be only weakly connected in childhood and adolescence*.

To fully capture the combined predictive potential of biological age measures on lifelong health, researchers might want to consider multivariate (e.g., including epigenetic and brain ages into one model as done in ^39^ and ^36^), and multimodal^40^ (e.g., structural, functional, diffusion MRI or different epigenetic markers not limited to DNAm) approaches.

### Strengths and limitations

This is the largest study that analyzed epigenetic and neuroimaging data from 15 international cohorts, often with repeated measures, across development (birth to early adulthood). We included a comprehensive selection of 28 epigenetic and brain age measures, a thorough consideration of potentially influencing factors, and a fully searchable toolbox (https://epi-brain-age-dashboard.streamlit.app/) to inform future studies.

Here, we focused on the error in chronological age prediction, a widely used benchmark, as an indicator for model performance. However, other benchmarks (e.g., association with age-relevant disease traits) might be equally or more relevant for health-related research questions^41,42^. Moreover, we were unable to derive brain ages in infancy and early adulthood was represented by a single dataset in some analyses. Furthermore, the observed small epigenetic PAR – brain PAR associations must also be interpreted considering measurement precision. As pooled prediction errors for both epigenetic and brain age exceeded the age range and correlations with chronological age were modest, the use of residuals of these moderately precise predictions may limit their reliability with downstream effects on association strength.

## Conclusion

With growing recognition that aging-related processes begin early in utero, evaluating how biological age measures perform during development is essential for studies of (healthy) aging across the life course. We found a dynamic developmental system of epigenetic-brain age performances and associations. Our study highlights the need for methodological advancements (including next-generation and longitudinal models) that focus on early development, laying the groundwork for future investigations into the lifespan trajectories of healthy aging.

## Data availability

The MIND consortium integrates data from multiple independent cohort studies. The individual-level data cannot be made publicly available because they are subject to the ethical approvals, informed consent provisions, and governance policies of the contributing cohorts. Researchers interested in accessing the underlying data should apply directly to the relevant cohort(s) in accordance with their established data access procedures. For further information or to collaborate as part of the MIND consortium, researchers can contact the MIND consortium at.

The informed consent obtained from ALSPAC (Avon Longitudinal Study of Parents and Children) participants does not allow the data to be made available through any third party maintained public repository. Supporting data are available from ALSPAC on request under the approved proposal number, B3067. Full instructions for applying for data access can be found here: http://www.bristol.ac.uk/alspac/researchers/access/. The ALSPAC study website contains details of all available data (http://www.bristol.ac.uk/alspac/researchers/our-data/).

## Funding and acknowledgments

We thank all participants (and their families) for taking part in the studies that are included in this project. We also thank all researchers and support staff at the individual research sites for making this project possible by contributing their data. We thank Prof Dan Stein, who passed away during this study, for his immense contributions to psychiatry research and team science in South Africa and across the world.

The secondary analyses reported in this manuscript have primarily been funded by UK Research and Innovation (UKRI) under the UK government’s Horizon Europe / ERC Frontier Research Guarantee [BrainHealth, grant number EP/Y015037/1 awarded to EW]. This funding supported EW, MS, and VB. Full acknowledgments for the MIND consortium cohorts and co-authors are outlined in the supplement. The funders of this study had no role in study design, data collection, data analysis, data interpretation, or writing of the report.

## Contribution statement

1. **Conceptualization (MIND project):** CAMC, EW, IKS, MS, and VB
2. **Methodology (MIND project):** CAMC, DJS, EPP, EW, IKS, JFF, JJT, JR, LWH, MB, MS, RLM, TPF, and VB
3. **Data acquisition (individual cohorts):** AMS, APT, CB, CH, CJW, CoM, CSM, DAN, DJS, DW, EBB, EPP, GS, HJZ, HK, HT, HVC, J-BP, JGE, JHP, JJT, JR, JS, JTN, KD, LK, LMR, LWH, MAM, MB, PDW, PF, PMP, RAB, RG-O, RLM, SAB, SB, SC, SE, TJS, TP, VKO, WL, and YYO
4. **Data analysis (individual cohorts):** AH, ALT, AMS, APT, CH, CJH, ClM, CoM, DAN, DC, EPP, EW, HP, IKS, JHP, JJT, JR, JRD, JS, KD, KJO, LMR, MAM, MB, MS, NSJ, PAR, PDW, RAB, RG-O, RT, SA, SB, SD, SH, SYC, TZ, VB, VC, and VNK
5. **Quality control (individual cohorts):** ALT, AMS, APT, CB, CH, CJH, CoM, DAN, DW, EPP, EW, HP, IKS, JFF, JHP, JJT, JR, JRD, JW, KS, LMR, MAM, MB, MS, NSJ, PAR, PF, PMP, RG-O, RT, SA, SAJ, SB, SH, SYC, TP, TPF, TZ, VC, and VNK
6. **Data curation and administration (individual cohorts):** AMS, APT, CH, CJH, CJW, CoM, DAN, DJS, DW, EPP, EW, GS, HJZ, HK, IKS, JFF, JGE, JHP, JJT, JR, JS, LMR, LWH, MAM, MB, MS, NK, PF, PMP, RAB, RG-O, RLM, SAB, SAJ, SB, SC, SD, SH, TJS, TP, TPF, VKO, VNK, and YYO
7. **Visualization:** EW, MS, and VB
8. **Writing – original draft:** MS and EW
9. **Writing – review & editing:** All authors - AH, ALT, AMS, APT, CAMC, CB, CDT, CH, CJH, CJW, ClM, CoM, CSM, DAN, DC, DJS, DW, EBB, EPP, EW, GS, HJZ, HK, HP, HT, HVC, IKS, J-BP, JFF, JGE, JHP, JJT, JR, JRD, JS, JTN, JW, KD, KJO, KS, LK, LMR, LWH, MAM, MB, MS, NK, NSJ, PAR, PDW, PF, PMP, RAB, RG-O, RLM, RT, SA, SAB, SAJ, SB, SC, SD, SE, SH, SYC, TJS, TP, TPF, TZ, VB, VC, VKO, VNK, WL, and YYO

