## Supplementary Information for "Epigenetic and brain age across development: Performance and associations in the MIND consortium"

### Table of contents

**Cohort and Co-Author Acknowledgments and Funding** 2
**Supplemental Methods** 6
SM 1.1 Discrepancies from the pre-registration 6
SM 1.2 Cohort information and information and in- and exclusion criteria 7
SM 1.3 DNA methylation preprocessing and excluded epigenetic age clocks 13
SM 1.4 MRI preprocessing and excluded brain age models 14
SM 1.5 Performance indicators 16
SM 1.6 Gestational age transformation 17
SM 1.7 Statistical analysis 17
**Supplemental Results** 22
SM 2.1 Performance of epigenetic clocks across development 22
SM 2.2 Epigenetic age performance moderators: Developmental period 23
SM 2.3 Epigenetic age performance moderators: Age range overlap between epigenetic training and testing data 25
SM 2.4 Epigenetic age performance moderators: Tissue overlap between epigenetic training and testing data 26
SM 2.5 Performance of brain age models across development 29
SM 2.6 Brain age performance moderators: Developmental period 30
SM 2.7 Brain age performance moderators: Age range overlap between brain age training and testing data 31
SM 2.8 Brain age performance moderators: Processing level (minimal vs extracted) 32
SM 2.9 Within-modality associations 33
SM 2.10 The influence of epigenetic clock generation on brain PAR - epigenetic PAR associations 36
SM 2.11 The influence of tissue on brain PAR - epigenetic PAR associations 37
SM 2.12 Sensitivity analyses of brain PAR–epigenetic PAR associations under alternative covariate adjustment 38
SM 2.13 Brain PAR - epigenetic PAR associations in a restricted subsample 39
**Supplemental references** 39
**References for all included biological age models** 42

**Cohort and Co-Author Acknowledgments and Funding**

**Individual**

**E.W., M.S.** and **V.B.** disclose support for the research of this work from UK Research and Innovation under the UK government's Horizon Europe / ERC Frontier Research Guarantee [BrainHealth, EP/Y015037/1].

Several authors disclose support for the research from by the European Union's HorizonEurope Research and Innovation Programme (FAMILY, grant agreement No 101057529 [**C.A.M.C, R.L.M, I.K.S**]; HappyMums, grant agreement No 101057390 [**C.A.M.C, I.K.S, R.G-O, L.M.R, V.K.O**]) and the European Research Council (TEMPO; grant agreement No 101039672 [**C.A.M.C, I.K.S, S.D**]). This research was conducted while **C.A.M.C** was a Hevolution/AFAR New Investigator Awardee in Aging Biology and Geroscience Research. Views and opinions expressed are however those of the author(s) only and do not necessarily reflect those of the European Union. Neither the European Union nor the granting authority can be held responsible for them.

**S.A.** discloses support for the research of this work from the National Institute of Environmental Health Sciences (T32ES012870 and F31ES037540).

**S.Y.C.** discloses support for the research of this work from the NMRC Open Fund – Young Individual Research Grant (MOH-001149-00).

**D.A.N.** discloses support for the research of this work from the National Institutes of Health (R01HD076592).

**J.J.T.** discloses support for the research of this work from the Juho Vainio Foundation, the Hospital District of Southwest Finland (Finnish State Grants for Clinical Research, ERVA/VTR), the Emil Aaltonen Foundation, the Alfred Kordelin Foundation, the Sigrid Jusélius Foundation, the Signe and Ane Gyllenberg Foundation and the Orion Research Foundation.

**C.J.W.** discloses support for the research of this work from the Science for Africa Foundation (Del-22-002), with support from the Wellcome Trust and the UK Foreign, Commonwealth & Development Office, as part of the EDCTP2 programme supported by the European Union.

**D.W.** discloses support for the research of this work from the Academy of Medical Sciences (Professorship APR7_1002). **T.P.F.** discloses support for the research of this work from a UKRI Future Leaders Fellowship (MR/Y017560/1).

**C.S.M.** discloses support for the research of this work from the National Institute of Mental Health (R01 MH103761, R01 MH130158, R01 MH121079), the Eunice Kennedy Shriver National Institute of Child Health and Human Development (R01 HD076592, R01 HD036916, U01HD110063), National Institute on Minority Health and Health Disparities (R01 MD011716), National Institutes on Aging (R01AG071071). The content is solely the responsibility of the authors and does not necessarily represent the official views of the National Institutes of Health.

**S.H.** discloses support for the research of this work from a GW4 BIOMED2 MRC/UKRI Doctoral Training Partnership studentship (MR/W006308/1).

**A.M.S.** discloses support for the research of this work from the National Center for Advancing Translational Sciences of the National Institutes of Health (TL1TR002371) and the Medical Research Foundation at Oregon Health & Science University (award #005137).

**Cohort**

***Avon Longitudinal Study of Parents and Children***

The UK Medical Research Council and Wellcome (Grant ref: MR/Z505924/1) and the University of Bristol provide core support for ALSPAC. This publication is the work of the authors and MS and EW will serve as guarantors for the contents of this paper. A comprehensive list of grants funding is available on the ALSPAC website (http://www.bristol.ac.uk/alspac/external/documents/grant-acknowledgements.pdf). Genomewide genotyping data was generated by Sample Logistics and Genotyping Facilities at Wellcome Sanger Institute and LabCorp (Laboratory Corporation of America) using support from 23andMe.

We are extremely grateful to all the families who took part in this study, the midwives for their help in recruiting them, and the whole ALSPAC team, which includes data collection staff, data and administrations staff, technical managers and the technical staff with the Bristol Bioresource Laboratory, based within the University of Bristol.

***Brazilian High Risk Cohort***

The Brazilian High Risk Cohort study was supported by the National Institute of Developmental Psychiatry for Children and Adolescents (INPD) with grants from: Fundação de Amparo à Pesquisa do Estado de São Paulo (FAPESP; 2008/57896-8, 2014/50917-0, 2013/08531-5, 2020/06172-1, 2021/12901-9, 2021/05332-8, 2023/00437-1, 2023/05560-6); Conselho Nacional de Desenvolvimento Científico e Tecnológico (CNPq; 573974/2008-0, 465550/2014-2); the European Research Council (337673 and 101057390); the UK Medical Research Council (MR/R022763/1); Ministério da Saúde (888379/2019 – Portaria no. 1.949, 04/08/2020); Banco Industrial do Brasil S/A; the CISM grant; and Coordenação de Aperfeiçoamento de Pessoal de Nível Superior (CAPES; code 001). Collaboration between the BHRC and other cohorts was funded by the National Institutes of Health (NIMH R01MH120482-01). Involvement of NIMH Intramural investigators was funded by the NIMH Intramural Research Program (Project MH 002782).

***Drakenstein Child Health Study***

The Drakenstein Child Health Study is supported by the Gates Foundation, the South African National Research Foundation (NRF), the Wellcome Trust, the South African Medical Research Council and the National Institutes of Health.

***Future of Families and Child Wellbeing Study /* Study of Adolescent to Adult Neurodevelopment**

The Future of Families and Child Wellbeing Study was supported by the Eunice Kennedy Shriver National Institute of Child Health and Human Development (R01-HD36916, R01-HD39135, U01-HD110063 and R01-HD40421) and a consortium of private foundations.

The Study of Adolescent to Adult Neurodevelopment (SAND) was supported by the US National Institutes of Health [R01 MH103761 (C.S.M), R01 HD076592 (Notterman), R01 MD011716 (Mitchell), R01 HD036916 (Edin & Waldfogel), R01 MH121079 (L.W.H, Mitchell, C.S.M)].

***Growing Up in Singapore Towards Healthy Outcomes (GUSTO)***

The GUSTO study is supported by the National Research Foundation (NRF) under the Open Fund–Large Collaborative Grant (OF-LCG; MOH-000504), administered by the Singapore Ministry of Health's National Medical Research Council (NMRC) and the Agency for Science, Technology and Research (A*STAR). Under RIE2025, the study is supported by funding from the NRF's Human Health and Potential (HHP) Domain, under the Human Potential Programme. Additional support was provided by the A*STAR Early Childhood Grant (H24P2M0005).

***Kids2Health***

The Kids2Health study was funded by the German Federal Ministry of Research, Technology and Space: 01GL1743 A/B (to C.H.) and 01GL1743-C (to E.B.), and 01KR1301-A and 01KR1301-B (to C.H.).

***Michigan Twin Neurogenetic Study***

This research was supported by the US National Institutes of Health (R01MH081813 and R01HD066040 to S.A.B.; R01HD104297 to S.A.B. and S.L.C.; R01HD093334 and UH3MH114249 to S.A.B. and L.W.H.). Additional support was provided by the Avielle Foundation (Conway Family Award for Excellence in Neuroscience, to L.W.H. and S.A.B.) and a NARSAD Young Investigator Grant from the Brain and Behavior Research Foundation (to L.W.H.), with institutional funding from the University of Michigan (to L.W.H.) and Michigan State University (to S.A.B.).

***Oregon ADHD-1000***

The Oregon ADHD-1000 cohort is supported by the National Institute of Mental Health (R01MH131685 to M.A.M. and J.T.N.; R37MH059105, R01MH115357 and R01MH099064 to J.T.N.) and by philanthropic support from the Steven J. Sharp Center for Mental Health Innovation at Oregon Health & Science University. Genomic DNA isolations were performed by the OHSU Integrated Genomics Laboratory (RRID: SCR_022651).

***The CannTeen Study***

The CannTeen cohort was supported by a grant from the UK Medical Research Council (MR/P012728/1) to H.V.C. and T.P.F.

#### *The FinnBrain Birth Cohort Study*

Hasse Karlsson was supported by Finnish State Grants for Clinical Research (ERVA), Signe and Ane Gyllenberg Foundation, and Research Council of Finland (# 314390, 332444). Linnea Karlsson was supported by Finnish State Grants for Clinical Research (ERVA), Signe and Ane Gyllenberg Foundation. Jetro Tuulari was supported by Finnish State Grants for Clinical Research (ERVA), Signe and Ane Gyllenberg Foundation, Alfred Kordelin Foundation, Juho Vainio Foundation, and Orion Research Foundation.

#### *The Generation R Study*

The Generation R Study is conducted by Erasmus MC, University Medical Center Rotterdam in close collaboration with the School of Law and Faculty of Social Sciences of the Erasmus University Rotterdam, the Municipal Health Service Rotterdam area, the Rotterdam Homecare Foundation and the Stichting Trombosedienst & Artsenlaboratorium Rijnmond (STAR-MDC), Rotterdam. The generation and management of the methylation data was executed by the Human Genotyping Facility of the Genetic Laboratory and the Genomics Core Facility of the Department of Internal Medicine, Erasmus MC. We gratefully acknowledge the contribution of children and parents, general practitioners, hospitals, midwives and pharmacies in Rotterdam. This project received funding from the European Union's Horizon Europe Research and Innovation Programme (STAGE, grant agreement No 101137146). The work of Charlotte Cecil, Ryan Muetzel and Isabel Schuurmans is supported by the European Union's HorizonEurope Research and Innovation Programme (FAMILY, grant agreement No 101057529 [CC, RM, IS]; HappyMums, grant agreement No 101057390 [CC, IS]) and the European Research Council (TEMPO; grant agreement No 101039672 [CC, IS]). This research was conducted while Charlotte Cecil was a Hevolution/AFAR New Investigator Awardee in Aging Biology and Geroscience Research. Views and opinions expressed are however those of the author(s) only and do not necessarily reflect those of the European Union. Neither the European Union nor the granting authority can be held responsible for them.

#### *The Neuroimaging of the Children's Attention Project*

This work was funded by the National Health and Medical Research Council of Australia (NHMRC; grant #2029361) and a grant from the Waterloo Foundation. The broader Children's Attention Project (CAP) and NICAP cohort was funded by NHMRC (grants #1065895 and #1008522). This work was supported by the MASSIVE HPC facility (www.massive.org.au). We thank all the children and families for their participation.

#### *Transitions in Adolescent Girls (TAG)*

Research reported here was supported by the National Institute of Mental Health (R01/R56 MH107418 and R01 MH127408; PI J.H.P.). The content is solely the responsibility of the authors and does not necessarily represent the official views of the National Institutes of Health.

#### *UC Irvine Daily Experiences in Pregnancy Study*

This work was funded by an ERANet Neuron grant [01EW1407A] and NIH R01HD-060628 and R01MD-017387.

### Supplemental Methods

#### SM 1.1 Discrepancies from the pre-registration

- *Cohorts*: Some cohorts in the pre-registration could ultimately not be included because i) sample size was insufficient after application of inclusion criteria and quality control or ii) the project timeline did not align with available resources. One additional cohort (TAG) joined the consortium and this project after pre-registration.
- *Biological age models:*
  - We now include both the EN and BLUP epigenetic age model by Zhang and colleagues[1].
  - The brain age model BabyPy was excluded as the underlying preprint was withdrawn. A new model and publication was not yet available when the current analyses were performed.
- *Modelwise meta-analysis approach*: In the pre-registration, we stated that the global meta-analyses (i.e., of epigenetic clock performance, brain age model performance, and brain age - epigenetic age associations) would be followed by model-specific meta-analyses (i.e., for individual epigenetic and brain age models and specific brain age - epigenetic age model combinations). This approach was changed to instead include model (or model combination) as a fixed effect in the global meta-analysis. This has the advantage that it uses one common cohort/timepoint variance structure and estimates all model means in the same model. That makes model estimates more directly comparable and avoids fitting many small meta-analyses with unstable heterogeneity estimates.
- *Associations and tissue*: We pre-registered that we would investigate the moderating influence of epigenetic tissue, to assess whether involvement of saliva tissue or cortical/brain tissue was associated with stronger brain age - epigenetic age associations. In the current analyses, we extended this to also include buccal tissue due to evidence suggesting that buccal tissue is more closely related to CNS tissue[2].
- *Additional analyses* (added based on discussions of the initial results and co-author feedback):
  - Meta-analyses of the unweighted mean absolute error (mainly reported in the supplement)
  - Moderation of epigenetic clock performance by clock generation
  - Exploration of whether age effects on epigenetic clock performance differ by epigenetic clock generation
- *Function to adjust postnatal age for gestational age*: There was a mistake in the pre-registration for this function, which has been corrected (see SM section 1.6).
- *Changes in False Discovery Rate (FDR) correction*: In the pre-registration we mistakenly stated that modelwise age effects would be adjusted across 28 (i.e., across modality). We now perform this adjustment by modality (i.e., across 17 for epigenetic clocks due to the exclusion of gestational clocks from these analyses and across 8 for brain age models). The significance of these individual analyses is not a focus of the current paper.
- *Pre-registered analyses that were not performed due to space constraints*:
  - Pearson's and Spearman's rank correlations between chronological age and biological age residuals (PAR) or biological age difference (PAD) scores

chronological age and between biological age difference scores (PAD) and biological age residuals (PAR)

- Performing meta-analyses of the weighted mean absolute error ($MAE_w$) without clocks that have not been trained to predict chronological age (PCGrimAge, PhenoAge, DNAmTL, DunedinPACE, DunedinPACNI). While DNAmTL and DunedinPACE, and DunedinPACNI were excluded from all analyses pertaining to the mean absolute error as this measure is not meaningful for these clocks, we refrained from rerunning analyses excluding the other next-generation clocks and instead considered the moderating influence of generation.
- Moderator analyses:
  - Test sample age range (brain & epigenetic performance + association)
  - Type of machine learning model (brain performance)
  - Mean age difference between MRI and DNA methylation (DNAm) assessments (associations)

#### SM 1.2 Cohort information and information and in- and exclusion criteria

##### Cohort information

Cohort information presented below was mostly extracted from the supplementary material of the MIND consortium paper[3], except for the TAG cohort, which joined the consortium after publication. Please also see SM Table S1 for an overview of all cohorts and key cohort overview papers.

***Avon Longitudinal Study of Parents and Children***

The Avon Longitudinal Study of Parents and Children (ALSPAC) study is a prospective birth cohort, which invited pregnant women resident in Avon, UK with expected dates of delivery between 1st April 1991 and 31st December 1992[4–6]. The initial number of pregnancies enrolled was 14,541, of which 13,988 children who were alive at 1 year of age. When the oldest children were approximately age 7, efforts were made to bolster the sample size. The total sample size for analyses using any data collected after the age of seven is therefore 15,447 pregnancies, resulting in 15,658 foetuses. Of these 14,901 children were alive at 1 year of age. Participants with DNAm data were derived from the Accessible Resource for Integrated Epigenomic Studies (ARIES[7]), which initially included a subset of 1,018 mother-offspring pairs and was later extended.

The present study included sub-samples of ALSPAC children with DNAm data at three timepoints: birth (n=891, 450k), age 7 (n=961, 450k), and age 17 (n=950, 450k; n=1,841, EPIC), and structural MRI data at age 20 (n=802). MRI-DNAm overlap sub-samples were n=172 (450k; MRI age 20.06 years, DNAm age 17.34 years) and n=305 (EPIC; MRI age 20.20 years, DNAm age 17.71 years).

Ethical approval for the study was obtained from the ALSPAC Ethics and Law Committee and the Local Research Ethics Committees. Informed consent for the use of all data collected was obtained from participants following the recommendations of the ALSPAC Ethics and Law Committee at the time. Participants can contact the study team at any time to retrospectively withdraw consent for their data to be used. Study participation is voluntary and during all data collection sweeps, information was provided on the intended use of data. The completion of a questionnaire, either on paper or online, was considered to be written consent from participants to use their data for research purposes. For the majority of tests undertaken during face to face visits, verbal consent was obtained from participants (both parents and children as appropriate) prior to the start of any data collection. However, some tests required the completion of a written consent form.

Biological samples are collected in accordance with the Human Tissue Act (2004). Specific Research Ethics Committee approval is sought for the consenting process at each collection sweep. Written consent, including permission for future use, is obtained from adult participants or from the parents of children as appropriate. Ethical approval for future use is covered by ALSPAC's Research Tissue Bank approval. All historical consents to hold biological samples have been reviewed as part of the Tissue Bank approval process. Participants can contact the study team at any time to retrospectively withdraw consent for use of their samples.

Please note that the study website contains details of all the data that is available through a fully searchable data dictionary and variable search tool:
http://www.bristol.ac.uk/alspac/researchers/our-data/.

***Brazilian High Risk Cohort - Happy Mums***

The Brazilian High-Risk Cohort is a prospective high-risk cohort that commenced in 2009-2010 and has been tracking children residing in Porto Alegre and São Paulo (Brazil) from mid-childhood onward. We used a two-stage design. We first assessed childhood symptoms and family history of psychiatric disorders in a screening interview, collecting information from 9,937 index children. In the second stage, a random subsample (intended to be representative of the community, n = 958) and a high-risk subsample (children at increased risk for mental disorders, based on family risk and childhood symptoms, n = 1554) were selected for further evaluation. From those, 750 children were invited to take part in a neuroimaging study and to provide blood samples for the assessment of peripheral blood biomarkers. These children, accompanied by their biological parents, participated in multiple research waves. The Scientific Committee and Research Ethical Commission of the Federal University of Rio Grande do Sul, and the Brazilian National Research Ethics Commission (CONEP), approved the protocol and informed consent form in accordance with the Brazilian and international regulatory framework under registration numbers IRB Nº: 20180018 and CAAE: 74563817.7.1001.5327.

***Drakenstein Child Health Study***

The DCHS is located in the periurban Drakenstein district, Western Cape, South Africa. Pregnant women were recruited while attending routine antenatal care at Mbekweni or TC Newman clinic between March 2012 and March 2015[8–10]. Pregnant mothers were eligible for the study if they were 18 years or older, planned to attend antenatal care at one of the two clinics and intended to remain in the area for at least a year. Expecting mothers provided informed written consent at enrolment and were reconsented annually following childbirth. In total, 1137 mother–child dyads were enrolled in the study, of which four mothers had twins and one had triplets. Thus, 1143 children were enrolled in the study. The study was approved by the faculty of Health Sciences, Human Research Ethics Committee, University of Cape Town, Stellenbosch University and the Western Cape Provincial Health Research committee.

***Future of Families and Child Wellbeing Study***

The Future of Families and Child Wellbeing Study (FFCWS) is based on a stratified, multistage sample of 4,898 children born in large U.S. cities (population over 200,000) between 1998 and 2000, where births to unmarried mothers were oversampled by a ratio of 3 to 1. This sampling strategy resulted in the inclusion of a large number of Black, Hispanic, and low-income families. Mothers were interviewed shortly after birth and fathers were interviewed at the hospital or by phone. Follow-up interviews were conducted when children were approximately ages 1, 3, 5, 9, 15, and 22 (still being collected). When weighted, the data are representative of births in large US cities.

***Growing Up in Singapore Towards Healthy Outcomes (GUSTO)***

Growing Up in Singapore Towards Healthy Outcomes (GUSTO) is a large longitudinal birth cohort study[11]. The GUSTO study recruited pregnant women aged 18 years and above, attending their first trimester antenatal dating ultrasound scan clinic at Singapore’s two major public maternity units, National University Hospital (NUH) and KK Women's and Children's Hospital (KKH), between June 2009 and September 2010. Participants were (i) Singapore citizens or permanent residents who were of (ii) Chinese, Malay or Indian ethnicity with homogeneous parental ethnic background, who (iii) had the intention of eventually delivering in NUH or KKH and (iv) intended to reside in Singapore for the next 5 years. Furthermore, (v) only women who agreed to donate birth tissues (including cord, placenta, and cord blood) at delivery were included. Mothers receiving chemotherapy, psychotropic drugs, or who had type I diabetes mellitus were excluded. A total of 1450 mothers were enrolled and 18 additional infants of low birthweight were enrolled at the delivery timepoint. Mother-child dyads were followed throughout pregnancy and beyond. The GUSTO study was approved by the National Healthcare Group Domain Specific Review Board (NHG DSRB) and the SingHealth Centralized Institutional Review Board (CIRB). Written consent was obtained from all guardians on behalf of the children enrolled in this study.

***Kids2Health – childhood and infancy***

The Kids2Health-Childhood study is a cohort of 535 children between 3-12 years of age in Berlin, Germany that was enriched for children with different types of chronic stress experiences (maltreatment, refugee stress, obesity as a metabolic stressor). Children were recruited between 2018 and 2023. The general design, research aims, and specific measurements of Kids2Health study have been approved by the Medical Ethical Committee of the Ludwig-Maximilians-Universität München. Written informed consent was obtained from the parents on behalf of the child. Kids2Health – infancy is a cohort of 181 infants between 0-2 years of age in Berlin, Germany that was enriched for children whose mothers were exposed to childhood maltreatment. Pregnant mothers were recruited in early pregnancy and underwent a detailed clinical interview regarding childhood adverse experiences and were then followed up three times during pregnancy, at delivery and study visits with their offspring took place at 1, 6, 12 and 24 months age. Recruitment was between 2018 and 2023. The general design, research aims, and specific measurements of Kids2Health study have been approved by the Medical Ethical Committee of the Ludwig-Maximililans-Universität München. Written informed consent was obtained from the parents on behalf of the child.

***Michigan Twin Neurogenetic Study***

Participants in the Michigan Twins Neurogenetics Study (MTwiNS) were recruited between 2016-2024 from the Twin Study of Behavioral and Emotional Development in Children (TBED-C), part of the broader Michigan State University Twin Registry[12,13]. The TBED-C identified twin families with children aged 6–10 years using birth records, focusing on families residing within 120 miles of East Lansing, Michigan. This region encompasses urban areas such as Detroit, Flint, and Lansing, along with suburban and rural locales. The overall sample comprised a population-based arm of 528 twin families and an "under-resourced" arm of 502 twin families from neighborhoods with above-average poverty levels, defined as over 10.5% of families living below the poverty line at the study's inception[14]. Detailed recruitment procedures are documented in[14]. The MRI sample was randomly sampled from 702 eligible disadvantaged families. The response rate among eligible families was 90.2%, of which 80.4% agreed to participate. We ultimately ran 93.3% of interested families. Conducted in the USA, MTwiNS included typical neuroimaging exclusions, such as excluding individuals with metal in their bodies. The enrolment details for women and children are as previously mentioned. The study received ethics approval and informed consent from the Institutional Review Boards (IRB) at both Michigan State University and the University of Michigan.

***Oregon ADHD-1000***

The Oregon-ADHD-1000 is a community-recruited, longitudinal, case-control cohort of children (N=1483; age 7–11 years at baseline; 39% female) from northwest Oregon (USA) that is enriched for psychopathology. Details about study recruitment and data collection procedures have been published previously[15–17]. First study visits were between 2009 and 2015. The local Institutional Review Board approved the studies. Parents provided written informed consent; children provided written informed assent. All families completed a multi-informant, multi-method screening process to establish eligibility and diagnostic group assignment for ADHD, non-ADHD, as well as comorbid disorders. The present study includes N=496 youth with good quality MRI and DNAm data.

***Transitions in Adolescent Girls (TAG)***

The Transitions in Adolescent Girls (TAG) study includes up to N=224 through two recruitment drives from Lane County, Oregon, USA. The original sample (N=174 female adolescents aged 10.0-12.99 years at enrollment) launched in 2015, and a replenishment sample (N=46, aged 14.0-16.99 years at enrollment) launched in 2022. Recruitment letters were sent to families with children in grade 5 or 6 that were registered as female by the schools, and this was supplemented (particularly for the replenishment sample) with information from secure databases of families who registered their interest in our lab or department's research, recruitment flyers posted around the community or disseminated at community events, and through snow-balling efforts.

Participants follow a two-visit protocol that includes hormone assessments, psychosocial functioning, neuroimaging, and anthropometric measures approximately every 18 months for a total of six waves. Each wave includes collection of:

- two in-lab saliva samples, four home saliva samples (one per week), and one hair sample for the assessment of hormone levels, immune factors, and methylation;
- an MRI session including structural, diffusion, resting-state functional and task-based functional scans;

- the Kiddie Schedule for Affective Disorders and Schizophrenia (K-SADS), a diagnostic interview on current and lifetime mental health;
- production of a short self-narrative video; and
- measurement of height, weight, and waist circumference.

In addition, adolescents and their parents/guardians complete a number of surveys to report on the adolescent's pubertal development, mental health, social environment and life events; adolescents also report on various indices of self-perception and social-emotional functioning.

Parents/guardians gave written informed consent and adolescents < 18 assent to participate. As adolescents reach age 18, they also gave written informed consent. Ethics approval was received from the Institutional Review Board of the University of Oregon (#03232015.027).

Inclusion criteria at time of enrollment of the original sample:
- Female
- 10, 11, or 12 years old
- Fluent in English
- Normal or corrected-to-normal vision

Exclusion criteria at time of enrollment of the original sample:
- Diagnosed with a developmental disability
- Diagnosed with a psychotic disorder
- Diagnosed with a behavioral disorder, including autism
- Taking psychotropic medication other than stimulants
- MRI contraindications (as described on the Lewis Center for Neuroimaging screening materials, including claustrophobia and presence of ferromagnetic material in the body)
- Report or suspect being pregnant

***The CannTeen Study***

The CannTeen study is a purposively recruited longitudinal cohort of adolescents (initial age 16-17 years) and adults (initial age 26-29 years) who either use cannabis regularly (at least weekly) or do not, based in London (UK). The initial sample comprised 274 participants, with 76 adolescents who use cannabis (50% girls), 63 adolescent controls (51% girls), 71 adults who use cannabis (47% women), and 64 adult controls (52% women). Participant were recruited between 2017 and 2019. Exclusion criteria were: daily use of psychiatric medication, current treatment for a mental health disorder, personal history of a psychotic disorder, and regular use of any other illicit drug (apart from cannabis). Full cohort details can be found in Lawn et al.[18]. All participants gave a saliva sample for epigenetic analyses. 140 participants (n=35 in each group) completed MRI scans at baseline and one year follow-up. The study received ethical approval from the University College London (UCL) ethics committee. As all participants were over 16 years of age, all participants gave their own informed consent to participate.

***The FinnBrain Birth Cohort Study***

FinnBrain Birth Cohort study is a prospective population-based cohort with recruitment around the city of Turku in Southwest Finland and the Åland island. Initial recruitment included 3808 mother-child dyads, which were recruited between December 2011 and April 2015. Participants for the neonatal neuroimaging study were 97 healthy Finnish (Caucasian) neonates (57 males, 40 females) from 2 to 5 weeks of age, counted from the estimated due dat. All the infants were born full-term [between gestational weeks 37 and 42] and weighed more than 2500 g. The exclusion criteria for the infants were: occurrence of any perinatal complications with neurological consequences (e.g., hypoxia), scoring lower than 5 points in the 5 min Apgar, previously diagnosed CNS anomaly or an abnormal finding in a previous MRI scan. The families were contacted via telephone and eligibility for the study was assessed. After explaining the purpose and protocol of the study, a written informed consent was signed by the parents on behalf of each infant. The study was conducted according to the Declaration of Helsinki and was reviewed and approved by the Ethics Committee of the Hospital District of Southwest Finland (ETMK: 31/180/2011).

***The Generation R Study***

The Generation R Study is a prospective population-based cohort that follows children from fetal life onwards. Pregnant women living in the study area of Rotterdam, The Netherlands, with an expected delivery date between April 2002 and January 2006 were invited to participate[19]. A total of 9,778 mothers were enrolled. These mothers, their children and partners took part in several research waves. The general design, research aims, and specific measurements of The Generation R Study have been approved by the Medical Ethical Committee of Erasmus MC, in accordance with the Declaration of Helsinki of the World Medical Association. Written informed consent was obtained from the parents on behalf of the child. For data collected when the child was 12 years or older, the child also provided written informed consent.

***The Neuroimaging of the Children's Attention Project***

The Neuroimaging of the Children's Attention Project (NICAP) is a longitudinal cohort examining brain and cognitive development in children with and without ADHD, with up to 3 repeated neuroimaging assessments between the ages 9-14. Data were collected between 2014 and 2019. NICAP as an extension of the non-imaging longitudinal study the Children's Attention Project recruited from the age 6-8years. Full cohort protocol details can be found in Silk, Genc[20] and Sciberras, Efron[21]. It is a community-based sample initially recruited from 43 socio-economically diverse primary schools across Melbourne, Australia. Exclusion criteria were: intellectual disability; previous known serious medical, neurological or genetic condition; moderate-severe sensory impairments; and insufficient English to participate. Total sample size is 365. A total of 471 multimodal MRI scans were collected across three waves of data over five years. Being a community sample, females are well represented, representing roughly half the typical control sample and ~25% of the ADHD sample. This study was approved by The Royal Children's Hospital (RCH) Human Research Ethics Committee (HREC #34071), and parents or guardians gave informed consent while participating children assent.

***UCIrvine Daily Experiences in Pregnancy Study***

The UCI cohort is a prospective, longitudinal study of ethnically and socio-demographically diverse pregnant women receiving prenatal care at the university and affiliated clinics, based in Irvine, California. Recruitment was between 2011 and 2014. All participants had singleton, intrauterine pregnancies, with no known cord, placental, or uterine anomalies, fetal congenital malformations, or presence of any conditions known to be associated with dysregulated neuroendocrine function or systemic corticosteroid medication use. The cohort comprised N = 253 mother-child dyads. All study procedures were approved by the university's IRB, and all participants (pregnant women, and parents on behalf of their infants) provided written informed consent.

**Inclusion and exclusion criteria of the current meta-analytical study**
The current study included cohorts that have both structural MRI and DNAm data available for at least one time point in participants up to the age of 24 (due to the developmental focus of the study). For three cohorts (FinnBrain, Kids2Health Infancy, UCI Echo) this criterion was relaxed because overlapping data were only available at birth, and age-specific preprocessing pipelines produced different imaging features that prevented application of the brain age models within the scope of the project. These cohorts were nevertheless included to acknowledge their contributions to the consortium and this project, despite only contributing a single modality per time point.

Each of the included cohorts has applied their own inclusion and exclusion criteria, including those that are standard in the field (e.g., MRI exclusion criteria such as pregnancy, and metal in the body, exclusion of participants with neurological disorders and a history of head trauma). Some cohorts applied additional inclusion criteria (e.g., ADHD-1000, which is an ADHD case-control study). These have been outlined in the MIND overview paper[3] and above.

For the current study, participants with poor MRI or DNAm quality were excluded. We also excluded samples (/timepoints within samples) that had less than 40 participants available for the association analyses (please see the section on 'Statistical Power' in our preregistration; https://osf.io/6csm3/files/tx5ea). The following exclusion approach was applied regarding clinically-relevant phenotypes (e.g., pre-term birth or psychiatric diagnosis such as ADHD): 'Cases' (e.g., preterm birth or a diagnosis of ADHD) were not excluded if the cohort is community-/population-based and as a result might include a small number of such 'cases'. However, 'cases' were excluded from case-control cohorts. This decision was made to ensure a consistent application of inclusion/exclusion criteria given that different studies would have assessed different variables and to adopt a population-based approach overall.

For further cohort details, see Schuurmans et al.[3].

#### SM 1.3 DNA methylation preprocessing and excluded epigenetic age clocks

*DNA methylation preprocessing and age estimation*

Each cohort completed a harmonized DNAm preprocessing pipeline (https://github.com/MarleneSt/MIND_BrainAge-EpiAge). The DNAm levels obtained from blood (cord, whole), placental, buccal or saliva tissues were assessed using the Illumina Infinium HumanMethylation450K BeadChip assay (Illumina 450 K array) and Infinium MethylationEPIC (v1 and v2) BeadChip (Illumina, San Diego, CA, USA) across more than 485,000, 850,000 and 930,000 CpG sites, respectively. For most cohorts, functional normalization and quality control (QC) were carried out using *meffil*[22] and included QC checks for sex, background detection and number of beads per sample or CpG. Methylation level at each CpG site was indexed via beta values. Cell counts were generated using the following reference panels: 'combined cord blood' (in *meffil*) for cord blood, 'blood gse35069' (in *meffil*) for whole blood, uniLIFE[23] for both cord and whole blood, and 'saliva gse48472' (in *meffil*) or hepiDish two-step method[24] for saliva or buccal tissue. Please see SM Table S3 for information on pre-processing in individual cohorts and deviations from this pipeline .

CpGs and samples with over 30% missing values were removed. Remaining CpGs were ordered by chromosome and genomic position and imputed using K-nearest neighbour (KNN) using the *impute*[25,26] package in R (https://bioconductor.org/packages/impute). CpGs that were entirely missing in a sample (e.g., CpGs exclusive to the EPIC array in a cohort with 450k data) were not imputed at this stage. Some of the included epigenetic age clocks performed additional imputation processes, including for CpGs that were entirely missing, usually via KNN or mean imputation. Extreme values were winsorised using 3 x IQR (Q1 - 3 x IQR to Q3 + 3 x IQR).

We generated epigenetic ages based on 20 publicly available clocks (see SM Table S2). Where possible, all epigenetic clocks were applied across all available timepoints in each cohort (birth to young adulthood), except for gestational clocks, which were only applied to birth timepoints and when gestational age information was available. In line with recommended guidelines, for two epigenetic clocks (AltumAge and Horvath2013), BMIQ normalization of DNAm data was performed using the *meffonym* (https://github.com/perishky/meffonym) package in R prior to deriving the epigenetic age estimates.

*Excluded clocks*

The following epigenetic clocks were excluded (see main text for inclusion criteria):

- Dunedin-PoAm[27] was excluded as we included the next generation/development of this clock (i.e., Dunedin-PACE).
- epiTOC and epiTOC2 [28] were excluded as both models were trained on mitotic division, a very specific phenotype with limited relevance to the brain, and primarily aimed to predict cancer, a (mostly) non-brain related disease.
- Mortality Risk Score[29] was excluded due to the training sample age range not overlapping with the age range of the current study (50-75 years).
- The NEOAge post-menstrual and post-natal age clocks[30] were excluded as they were trained exclusively on very preterm infants.
- Pan-mammalian clocks 2/3[31] were excluded as they are cross-species clocks, which is not the focus of this project.

#### SM 1.4 MRI preprocessing and excluded brain age models

*MRI preprocessing*

For details on MRI preprocessing per cohort, please see SM Table S4.

Structural T1-weighted MRI scans were acquired using 1.5 or 3T scanners and preprocessed using FreeSurfer version 6 or higher[32]. All cohorts quality controlled their neuroimaging data using one or more provided QC options (e.g., MRIQC[33], Euler number[34], standard ENIGMA QC protocols [https://github.com/ENIGMA-git/ENIGMA-FreeSurfer-protocol/tree/main]).

Missingness primarily emerged for samples where a region-by-region quality control protocol is applied to the FreeSurfer pre-processed data (e.g., ENIGMA QC). Here, missing values were imputed prior to applying the brain age models using the *missForest* R package[35].

We generated brain ages based on 8 publicly available clocks (see SM Table S2). While all cohorts generally followed the MIND protocols for estimating brain ages (see Step 5 https://github.com/MarleneSt/MIND_BrainAge-EpiAge), we note the following deviations:

- Kaufmann brain age model in FFCWS and MTwiNS: While the MIND pipeline generates FS-style surface parcel outputs (focusing on the cortical ribbon) and explicitly writes the stats files that are used subsequently, the scripts used for these two cohorts generates a volumetric subject-space parcellation NIfTI. This difference might lead to slight differences in parcel definitions but due to the meta-analytical nature of the current project, this was not deemed a problem. Additionally, leave-one-cohort-out analyses were not concerning.
- DBN brain age model in TAG: In order to generate outputs successfully the script for predicting brain age based on the DBN model was amended. Specifically, the (minimal) pre-processing steps were amended whereby processing was performed outside of ANTsPyNet without N4 bias correction and affine registration to MNI space. Leave-one-cohort-out analyses indicated that results remained consistent when removing TAG from DBN modelling and the cohort was therefore retained in these analyses.

*Excluded brain age models*

Based on our inclusion criteria (see main text), the following brain age models were excluded:

- Abam[36] was excluded as it focused on overlapping features to multiple already included models (Desikan-Killiany extracted values) whilst being trained on a comparatively small sample (compared to the included models). Additionally, the shared pre-trained model was not the best performing model in Brouwer et al.'s study.
- brainageR[37] and MCCQR[38] were excluded as their processing involved the neuroimaging software SPM.
- CentileBrain BrainAge1[39] was excluded from the current project as we are including its further developed version, CentileBrain BrainAge2, which was trained on substantially more participants[40].
- fetalBA[41] was excluded as the brain age model was exclusively trained on in-utero fetuses which is not the focus of the current study.
- Neuroanatomical Age Prediction using R (NAPR[42]): As part of this model, FreeSurfer-derived cortical thickness maps are transferred to the OpenCPU server. When testing the workflow, we were unable to connect to the server and additionally

expect that due to privacy regulations, the use of Amazon Web Services would not be possible for several MIND cohorts.

- SPARE-BA (e.g.,[43]) was excluded as it included a pre-processing approach that went beyond minimal processing that did not rely on FreeSurfer (e.g., regional volumetric maps calculated with DRAMMS) and the model was trained on participants ages 48 years and older.
- VolBrain BrainStructureAges (BSA[44]) was excluded as it required the individual upload of T1 scans to a website, which we deemed to not be feasible for the current project for time and privacy reasons.
- BabyPy was excluded as the preprint was withdrawn after our study was preregistered.

#### SM 1.5 Performance indicators

In total, we assessed the following complementary performance indicators, which are reported on in the main manuscript ($MAE_w$, correlations), the SMs (unweighted MAE) as well online (all others; https://epi-brain-age-dashboard.streamlit.app/).

**Unweighted Mean Absolute Error (MAE)** between chronological and biological age with larger values indicating a larger disparity between chronological and biological age.

**Root Mean Squared Error (RMSE)**, i.e., the square root of the average of squared errors with lower values indicating better fit.

**MAE weighted by the test sample age range ($MAE_w$** = MAE/max age – min age test sample**)**: MAE with increased comparability between cohorts with differing age ranges, with larger values indicating a larger disparity between chronological and biological age.

**MAE weighted by the training sample age range ($wMAE_{training}$** = MAE/max age – min age training sample): MAE with increased comparability between models that have been trained on different data from varying age ranges, with larger values indicating a larger disparity between chronological and biological age.

**Normalised RMSE (nRMSE)**: RMSE that is less sensitive to the age range of the sample (nRMSE = RMSE/max age – min age test sample), with lower values indicating better fit.

**Relative Absolute Error (RAE):** absolute error that is less sensitive to the age range of the test sample, with lower values indicating better fit[45].

$$RAE = \frac{\Sigma_{i=0}^{N} | \hat{y_i} - y_i}{\Sigma_{i=0}^{N} | y_i - \underline{y}}$$

**Correlation** between brain/epigenetic age and chronological age (Pearson's correlation coefficient) with higher values indicating better fit.

**$R^2$**: the proportion of variance in chronological age that is explained by the predicted biological age, with higher values indicating better fit.

### SM 1.6 Gestational age transformation

To facilitate integration of age at birth timepoints and gestational clocks (which produce predicted age in gestational weeks) into subsequent meta-analyses, these were transformed into age in years relative to gestational age using the following formula[46].

For gestational age for birth timepoints and predicted gestational age from gestational epigenetic age clocks, the following formula was applied:

$$actual/predicted\ gestational\ age\ in\ years = 7 * (actual/predicted\ gestational\ weeks - 40) \div 365$$

For the youngest cohorts/timepoints (ALSPAC, Kids2Health Infancy, Generation R, UCI cohort), chronological age was adjusted for gestational age to account for differences in gestational periods / postmenstrual ages. The exception to this were the timepoints M3 and M9 in the GUSTO cohort, which although < 2 years, did not undergo this adjustment.

$$adjusted\ chronological\ age = chronological\ (postnatal)\ age\ in\ years + gestational\ age\ in\ years$$

For participants where gestational age at birth was not available, a gestational period of 40 weeks (0.77 years) was assumed. This adjustment was applied up to age two years by which pre-term infants were assumed to have caught up to some extent with their full-term peers[47,48] and the influence of gestational age should have attenuated. The adjustment was not applied for gestational epigenetic clocks as these are trained on gestational age.

### SM 1.7 Statistical analysis

*Meta-analyses*

We performed inverse-variance weighted multi-level random-effects meta-analyses using the *metafor* R package[49], accounting for nesting in cohorts and time points (and models), to determine (e.g., $I^2$) and explore (moderator analyses) heterogeneity and provide pooled effect estimates.

<u>Performance</u>

*Meta-analysis*

To assess heterogeneity in how different epigenetic and brain age models perform in predicting chronological age across development, we performed two separate meta-analyses across all brain age models and epigenetic clocks. The main meta-analyzed metric was MAEw. As the MAE may be less informative for clocks that have not been trained to predict chronological age (PCGrimAge, PhenoAge, DNAmTL, DunedinPACE, DunedinPACNI), we performed an additional meta-analysis focused on **Pearson's correlation coefficient.** Notably, DNAmTL, one of the epigenetic clocks, is an estimator of leukocyte telomere length measured in kilobases, whereby larger values (i.e., a longer telomere length) is associated with younger age. Therefore, correlations between DNAmTL and chronological age were inverted prior to inclusion in meta-analyses.

To examine which features might explain heterogeneity in model performance, the following moderators were explored:

- **Epigenetic & brain age models**
  - Brain age or epigenetic age model to acquire pooled performance estimates for individual models
  - Sample mean age (these analyses exclude gestational epigenetic clocks as they were only applied at birth)
  - Overlap of training and test sample age range (these analyses exclude gestational epigenetic clocks as they were only applied within age range)
- **Epigenetic age models**
  - Tissue (training sample, test sample, and their correspondence)
- **Brain age model**
  - Processing level (minimal vs extracted)

Additionally, we investigated the interaction between model and sample mean age to explore age effects at the individual model level.

Notably, for the epigenetic meta-analyses, the $MAE_w$ was log-transformed due to substantial differences in variances across effects. Reported results related to the $MAE_w$ are back transformed unless otherwise specified. Moderator effects reported on the log scale are referred to with $b_{log.}$ For meta-analyses of Pearson's correlation coefficient, coefficients were Fisher z transformed. Reported results related to Pearson's correlation are back transformed unless otherwise specified. Moderator effects reported on the Fisher z scale are referred to with $b_z$. Lastly, supplementary analyses were performed for the raw/unweighted MAE. Here, the unweighted MAE was log-transformed and a different optimizer (BFGS instead of the default nlminb) was applied to ensure convergence.

Associations

*Cohort-level analyses*

To assess brain age - epigenetic age associations, cross-sectional measures of brain and epigenetic age per cohort and time point were analyzed. We performed robust linear regression analyses (via the *rlm* function from the *MASS* R package) with heteroscedasticity consistent standard errors with the brain-predicted age residual (brain-PAR) as the outcome and epigenetic-predicted age residual (epigenetic-PAR) as the predictor. *rlm* uses M-estimators to down-weigh outliers and tends to be more robust against skewed distribution of residuals, heavy tails, heteroscedasticity, as well as small-to-moderate violations of normality[50]. PAR was chosen for both epigenetic and brain age as this measure is adjusted for age-dependency and allows for better comparisons across cohorts with different age structures[51].

**Model 1 (primary model):** brain-PAR ~ epigenetic-PAR + age at MRI + age at methylation + sex + batch

- While the use of PAR adjusts for age-dependency, residual confounding by age may persist including because biological aging mechanisms are linked to time[52]. Additionally, methodological research suggests that omitting age when using detrended outcomes such as PAR may bias effect estimates and precision[52,53]. Therefore, age was included as a covariate. For cohorts where DNAm and MRI were not assessed at the same time, both ages were included in the model unless they correlated > 0.8 in which case only age at MRI was included.

- We decided against including age$^2$ in these models due to the often narrow age ranges per time point in many of the samples, within which we would mostly expect linear age-related changes in brain structure and DNAm[54,55].

**Model 2 (uncorrected model – supplementary analysis):** brain-PAR ~ epigenetic-PAR

**Model 3 (extended model – supplementary analysis):** Model 1 with cell type as an additional covariate: brain-PAR ~ epigenetic-PAR + age at MRI + age at methylation + sex + batch + cell type

- Cell type is not included in Model 1 as the exact role of cell type in epigenetic age estimates is unclear. While it may function as a confounder[56], adjusting for cell type composition may also remove part of the biological ageing signal due to changes in cell proportion being associated with ageing. Hence, Model 1 and 2 did not adjust for cell type while Model 3 allowed us to assess the impact of cell type and whether previously observed associations between epigenetic and brain age are driven by confounding due to cell type shifts.

Overall, this resulted in 136 association analyses per cohort at each available time point for the main analyses regressing brain-PAR onto epigenetic-PAR for each of the three models outlined above (i.e., 8 brain age models associated with 17 epigenetic clocks as the 3 gestational clocks had no brain age model equivalent). Notably, association analyses were not run for birth timepoints as we only had epigenetic ages but not brain ages at this developmental timepoint. It is notable that some cohorts were not able to apply every epigenetic clock or brain age model (e.g., neonatal cohorts that have to undergo different pre-processing procedures, cohorts with high missingness for certain CpGs, data privacy concerns for models that require uploading data to a website). Hence, the final number of analyses per cohort varies.

In addition to the associations between brain age and epigenetic age measures, we also assessed the associations 1) between measures derived from different epigenetic clocks (136 analyses), and 2) between measures from different brain age models (28 analyses), using zero-order correlations.

*Meta-analysis*

Subsequently, we meta-analyzed the standardized betas derived from the cohort-level analyses for each timepoint and model combination using inverse-variance weighted multi-level random-effects meta-analyses, accounting for nesting in cohorts and time points and model combination as outlined above. Notably, as for the Pearson's correlation coefficient with chronological age, associations between DNAmTL and brain age residuals were inverted prior to inclusion in meta-analyses.

We ran a meta-analysis across all brain age-epigenetic age model combinations to assess heterogeneity. The following moderators were analyzed:

- Model combination (of which there were 136) to acquire pooled performance estimates for individual model combinations (reflecting the specific brain age model - epi age model pair).
- Sample mean age
- Trained phenotype (in relation to epigenetic age models)
- Tissue (in relation to epigenetic age models): cortical vs non-cortical, saliva vs non-saliva, buccal vs non-buccal.

Additionally, we investigated the interaction between model combination and sample mean age to explore age effects at the level of specific brain age and epigenetic age model pairs (e.g., Horvath - ENIGMA and AltumAge - Centile2).

Insights from the proposed meta-analyses, heterogeneity estimates, and moderator analyses were used to infer whether certain model combinations show stronger associations, which features influence the strength of association and how associations change across development.

**Inference criteria**

Throughout this project, emphasis was placed on effect sizes and heterogeneity estimates (meta-analysis) over significance.

<u>Cohort-level analyses</u>: As the focus was on effect sizes and subsequent meta-analyses, no multiple comparison corrections were applied to the cohort-level analyses.

<u>Meta-analysis</u>:

The performance meta-analyses focused on the MAE, whose significance itself is not of relevance. Instead, multiple comparison corrections focused on the moderator analyses that explore heterogeneity. For the first performance meta-analysis combining the MAEs of all brain age models and all epigenetic clocks, respectively, we adjusted the *p*-values for the individual moderator analyses for multiple comparisons using the FDR (see SM Table S1.7.1). In the meta-analyses where we derived model-specific estimates by including model as a fixed effect, we only explored one moderator (age/developmental period), and we adjusted across the number of individual meta-analyses (15 [MAEw] and 17 [*r*] for epigenetic age due to exclusion of gestational clocks in these analyses, 15 [MAEw] and 17 [*r*] for brain age) when assessing its significance.

For the meta-analyses of the associations (standardized beta) between all brain age-epigenetic age combinations, we applied an FDR correction across the moderator analyses (see SM Table S1.7.1). For the association meta-analyses focused on each individual brain age-epigenetic age model combination by including a fixed effect for model combination, we corrected the significance of the association across the 136 performed meta-analyses. Additionally, we explored one moderator (age/developmental period) and therefore adjusted across the number of moderator analyses (136) when assessing its significance.

**SM Table S1.7.1. FDR correction approach for the moderator analyses**

| Epigenetic age performance | Brain age performance | Associations |
|---|---|---|
| model | model | model combinations |
| age | age | age |
| age range overlap | age range overlap | generation (2 level) |
| tissue overlap | processing level | generation (4 level) |
| tissue test | | saliva test or training |
| tissue training | | buccal test or training |
| generation (4 level) | | cortex training |
| FDR correction across 7 moderators per considered outcome | FDR correction across 4 moderators per considered outcome | FDR correction across 6 moderators per considered outcome |

**Assumption Violation/Model Non-Convergence**

For the association analyses, we adopted the following approach: We ran an initial set of analyses in two samples (ALSPAC and Generation R), covering mid-childhood to late adolescence/early adulthood to assess the suitability of linear regression models. Specifically, we performed linear models (via the *lm* function in R) with heteroscedasticity consistent standard errors and generated diagnostic plots (i.e., residuals versus fitted values, Q-Q, spread-location, and residuals versus leverage plots). These were inspected across the two samples to assess violations of regression assumptions. As we observed violations and as preregistered, we shifted our analytical approach to robust linear regression models (via the *rlm* function from the *MASS* R package) with heteroscedasticity consistent standard errors as outlined above.

### Supplemental Results

#### SM 2.1 Performance of epigenetic clocks across development

*Unweighted MAE*

Across 667 epigenetic age model performance estimates (i.e., number of effects across clocks, cohorts, and timepoints), the pooled MAE was 6.61 years (95% CI: 2.78–15.70), with substantial between-model heterogeneity ($I^2$=100%), of which ~96% was attributable to differences between clocks. Leave-one-cohort-out analyses showed stable pooled estimates (MAE range: 6.14–6.79 years), indicating no single cohort disproportionately influenced results.

Clock performance varied markedly across models (QM(18) = 11046837.83, $p_{uncorrected} < .001$), with gestational clocks (e.g., Bohlin, Knight) showing the lowest MAEs and later-life clocks (e.g., DamAge, AdaptAge, PhenoAge) showing substantially larger prediction errors. Pooled MAEs ranged from 0.06 years for Bohlin to 69.50 years for DamAge. Heterogeneity remained extremely high ($I^2$=99.96%). Leave-one-cohort-out analyses demonstrated relatively robust model-specific estimates, with minimal fluctuation in MAE after exclusion of individual cohorts for most models. The largest differences were observed for PhenoAge (range= 19.99 - 34.15) and ZhangEN (range = 16.92 - 27.21), where the exclusion of GenR resulted in a much lower pooled MAE.

*Weighted MAE by generation*

Multilevel meta-analysis demonstrated substantial differences in $MAE_w$ across epigenetic clock generations (QM(4) = 93.96, $p_{FDR} < .001$). In the generation-only model, fourth-generation clocks showed the highest prediction error, followed by second-generation clocks, while first-generation clocks showed the lowest prediction error. Gestational clocks showed substantially lower prediction error than all other clock generations. Notably, CIs were wide, especially for second and fourth generation clocks. $MAE_w$ by generation were:

- 0.08 (95% CI 0.03–0.23) for first-generation gestational clocks
- 2.87 (95% CI 1.26–6.62) for first-generation clocks
- 7.69 (95% CI 2.27–26.08) for second-generation clocks
- 19.86 (95% CI 5.86–67.35) for fourth-generation clocks

It is important to note that gestational clocks were only applied to birth time points whereas all other clocks were applied across all ages (birth to young adulthood), including outside their training ages. Hence, superior performance of gestational clocks was expected, and their performance is not directly comparable to that of other clocks.

*Weighted MAE in a restricted subsample*

We reran the main performance ($MAE_w$) meta-analysis after restricting it to effect sizes derived from samples/timepoints where (1) the age ranges of the test data and training data overlapped completely and (2) the training and test data were derived from the same tissue. This subset included 29.1% of the effect sizes from the main $MAE_w$ analyses and retained 11 of the 19 clocks (61%), including all three gestational clocks.

As expected, because greater age and tissue overlap were associated with improved performance (see SM 2.3 and 2.4), the pooled $MAE_w$ was substantially lower in this restricted analysis (0.95, 95% CI [0.38, 2.36]) than in the full analysis (2.18). Heterogeneity nevertheless remained high ($I^2$ = 99.9%, $p$ < .001).

*Pearson correlations*

The overall pooled Pearson correlation between predicted and chronological age across all epigenetic clocks, cohorts and developmental periods was moderate in magnitude ($r$ = .35, 95% CI [.17, .50]), indicating that epigenetic age estimates showed a modest overall correspondence with chronological age across development. We observed substantial heterogeneity across estimates ($I^2$ = 98.53%, Q(734) = 23645.97, $p$ < .001). Variance decomposition indicated that heterogeneity was primarily attributable to differences between clock models (54.81%), followed by cohort-level differences (38.59%) and, to a lesser extent, developmental time points within cohorts (5.14%). Leave-one-cohort-out analyses indicated that the influence of individual cohorts on these findings was small, with pooled correlations remaining highly stable across cohort exclusions (pooled $r$ range = .32–.37).

Modelwise analyses further highlighted large differences in predictive performance between clocks (QM(20)= 6779.55, $p_{FDR}$ < .001). Gestational clocks demonstrated the strongest correspondence with chronological age, including Bohlin ($r$ = .81, 95% CI [.76, .85]), EPIC ($r$ = .78, 95% CI [.72, .83]) and Knight ($r$ = .63, 95% CI [.55, .70]). In contrast, next-generation clocks generally showed weaker associations with chronological age, including DNAmTL ($r$ = .16, 95% CI [.05, .28]) and DunedinPACE ($r$ = .10 95% CI [-.02, .23]). See SM Table S10 for detailed modelwise results.

*Pearson correlations by generation*

Multilevel meta-analysis demonstrated substantial differences in Pearson's correlation coefficients with age across epigenetic clock generations (QM(5) = 165.37, $p_{FDR}$ < .001). In the generation-only model, fourth-generation clocks showed the highest prediction error, followed by second-generation clocks, while first-generation clocks showed the lowest prediction error. Gestational clocks showed substantially lower prediction error than all other clock generations. Notably, CIs were wide, especially for second and fourth generation clocks. Pearson correlation coefficients by generation were:

- .75 (95% CI .67 - .82) for first-generation gestational clocks
- .29 (95% CI .16 - .41) for first-generation clocks
- .22 (95% CI .06 - .38) for second-generation clocks
- .10 (95% CI -.13 - .32) for third-generation clocks (DunedinPACE only)
- .16 (95% CI -.03 - .33) for fourth-generation clocks

**SM 2.2 Epigenetic age performance moderators: Developmental period**

*Weighted MAE by generation*

Across all clock generations (but excluding gestational clocks), prediction error significantly decreased with increasing sample age. When including a mean age by clock generation moderation term (QM(6) = 676.18, $p_{uncorrected}$ < .001), geometric mean $MAE_w$ decreased by (these analyses exclude gestational clocks as these were only estimated at birth/one age):

- 11.10% per year for first-generation clocks (95% CI -14.47% to -7.60%)
- 13.82% per year for second-generation clocks (95% CI -17.09% to -10.42%)
- 12.34% per year for fourth-generation clocks (95% CI -15.66% to -8.88%)

Interaction models using first-generation clocks as the reference group indicated significantly steeper age-related reductions in prediction error for second-generation clocks (*b* = -0.03, $p_{uncorrected}$ < .001) and fourth-generation clocks (*b* = -0.014, $p_{uncorrected}$ < .001). Although age-related improvements in prediction accuracy were observed across all generations, the magnitude of these improvements differed modestly between generations, with second-generation clocks showing the steepest age-related reductions in prediction error.

Leave-one-cohort-out analyses demonstrated robustness of the estimated age effects across cohorts. Estimated annual percentage changes in $MAE_w$ varied only modestly across leave-one-out iterations:

- -11.76% to -10.03% for first-generation clocks
- -14.87% to -13.01% for second-generation clocks
- -13.98% to -10.11% for fourth-generation clocks

Overall, these findings suggest that epigenetic clock generation is strongly associated with performance, while age-related improvements in prediction accuracy are broadly consistent but modestly steeper for later-generation clocks.

*Pearson correlations*

In contrast to $MAE_w$, the overall association between epigenetic age estimates and chronological age did not significantly change across childhood and adolescence ($b_z$ = 0.005, $p_{FDR}$ = .154), indicating that the pooled correlation between predicted and chronological age remained comparatively stable across development. Nevertheless, substantial heterogeneity remained (QE(df=715) = 18440.51, $p$ < .001, $I^2$ = 96.82%), with most variability attributable to cohorts (73.02%), followed by developmental time points within cohorts (12.01%) and clock models (11.79%).

Leave-one-cohort-out analyses indicated that these findings were stable across cohort exclusions ($b_z$ range = 0.002 - 0.007).

Despite the absence of a significant overall age effect, modelwise analyses revealed differences in age effects between clocks (SM Figure S1). Several clocks showed increasingly stronger correlations with chronological age in older samples, including cAge, ZhangEN, ZhangBLUB, skinHorvath, DNAmTL and PCGrimAge. In contrast, PCBrainAge, and DunedinPACE and showed significantly weaker correlations with chronological age in older cohorts. Other clocks, including AdaptAge, PedBE, CorticalClock, Hannum, Horvath2013 and PhenoAge, showed comparatively stable associations across development.

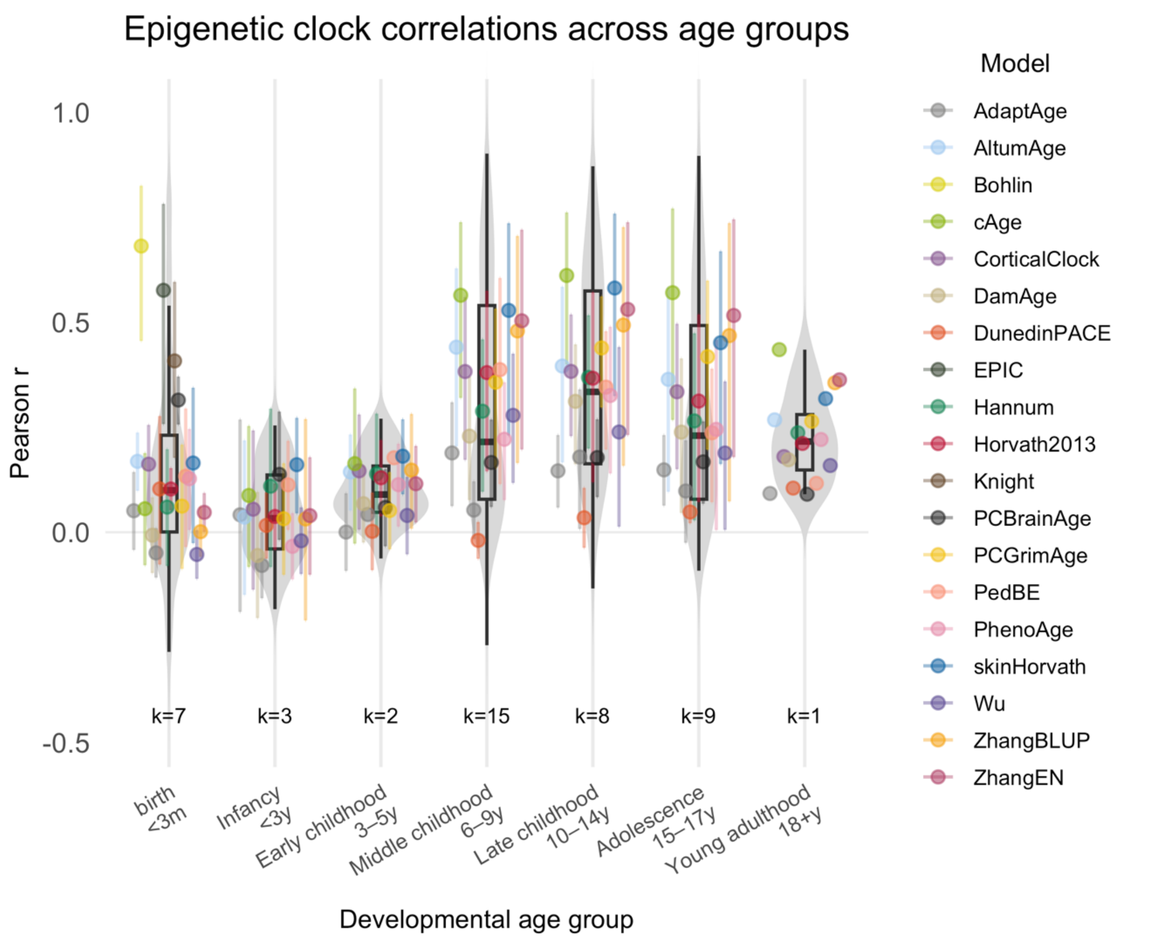


**SM Figure S1.** Epigenetic clock correlations across age groups.
k reflects the number of unique cohort-timepoint combinations contributing to the model-specific meta-analyses in each age bracket. Notably, if a cohort has multiple tissues or arrays at a timepoint, these are considered as two timepoints (e.g., 6years_blood and 6years_saliva).

Allowing age effects to vary across clocks significantly reduced heterogeneity (QM(34)=3551.37, $p_{uncorrected} < .001$), but overall heterogeneity remained high (QE(683) = 14785.11, $p < .001$, $I^2 = 96.24$). Leave-one-cohort-out analyses demonstrated robustness of the model-specific developmental effects, indicating no disproportionate influence of individual cohorts on the observed developmental patterns.

DunedinPACE ($\beta = -0.009$, $p_{FDR} = .029$) showed a significant negative age slope, reflecting decreasing correlations with chronological age in older developmental samples.

#### SM 2.3 Epigenetic age performance moderators: Age range overlap between epigenetic training and testing data

*Weighted MAE*

Age overlap was defined as the percentage of the test sample age range that is covered by the training age range (overlap / study age range).

A greater proportional overlap between the cohort age range and the epigenetic clock training age range was associated with significantly lower geometric mean $MAE_w$, with each 1% increase in overlap associated with an approximately 0.6% reduction in geometric mean $MAE_w$ ($b_{log}$ = -0.006 log units, 95% CI [-0.007,-0.006], $p_{FDR} < .001$). Predicted geometric mean $MAE_w$ decreased from approximately 5.95 at 0% overlap to 3.04 at 100% overlap, indicating progressively better apparent epigenetic age prediction performance with increasing training-age overlap. Proportional age overlap explained a substantial proportion of between-study heterogeneity (QM(1) = 1277.40, $p_{FDR} < .001$), and leave-one-cohort-out analyses showed that the effect was robust, remaining consistently negative across all re-fitted models; each 1% increase in overlap was associated with a reduction in geometric mean $MAE_w$ ranging from -0.72% to -0.59% across cohort exclusions (SD = 0.03%).

*Pearson correlations*

Greater proportional overlap between the epigenetic clock training-age range and the developmental test-sample age range was associated with significantly stronger correlations between predicted and chronological age ($b_z$ = 0.0017 Fisher-z units per 1% overlap increase, 95% CI [0.0016, 0.0019], $p_{FDR} < .001$). Predicted pooled correlations increased from $r = .18$ in samples with no overlap to $r = .35$ in samples with complete (100%) overlap between training and test age ranges.

Proportional age overlap explained a substantial proportion of between-study heterogeneity (QM(1) = 436.89, $p_{FDR} < .001$), although considerable residual heterogeneity remained (QE(715) = 16832.88, $p < .001$; $I^2 = 96.60\%$). Variance decomposition indicated that heterogeneity was primarily attributable to cohort differences (77.93% of observed variance), followed by developmental time points within cohorts (12.44%) and clock-model differences (6.22%).

Leave-one-cohort-out analyses demonstrated that the proportional overlap effect was highly robust and not driven by any single cohort: the overlap effect remained consistently positive across all re-fitted models ($b_z$ = 0.00160–0.00192 Fisher-z units per 1% overlap increase; SD = 0.00008).

#### SM 2.4 Epigenetic age performance moderators: Tissue overlap between epigenetic training and testing data

<u>*Weighted MAE*</u>

*Tissue overlap*

Tissue overlap was defined as the same training and test tissue. For multi tissue clocks all test tissues were considered to be overlapping. Epigenetic age models performed significantly better when the tissue type of the test sample overlapped with the tissue(s) used during model training, with tissue overlap associated with an approximately 22% lower geometric mean $MAE_w$ ($b_{log}$ = -0.22 log units, 95% CI [-0.24,-0.21], $p_{FDR} < .001$. Predicted geometric mean $MAE_w$ was 2.41 in non-matching samples versus 1.92 in overlapping samples. Tissue overlap explained a substantial proportion of between-study heterogeneity (QM(1) = 716.45, $p_{FDR}$ < .001), indicating that concordance between training and test tissue type is an important determinant of epigenetic age prediction performance. Leave-one-cohort-out analyses showed that the tissue-overlap effect was robust and consistently negative across all re-fitted models, with tissue overlap associated with reductions in geometric mean $MAE_w$ ranging from -12.27% to -25.28% across cohort exclusions (SD = 3.35%).

*Tissue of the training sample (cortex, blood, multi, buccal, cord)*

Training tissue was significantly associated with epigenetic age prediction performance (QM(5) = 60.24, $p_{FDR}$ < .001), with substantial differences in geometric mean $MAE_w$ observed across tissue-training groups. Cord-trained clocks showed the lowest predicted geometric mean $MAE_w$ (0.08, 95% CI [0.02, 0.26]), followed by buccal-trained (1.65, 95% CI [0.26, 10.69]) and multi-tissue-trained clocks (1.81, 95% CI [0.54, 6.14]), whereas blood-trained (5.54, 95% CI [2.20, 13.52]) and cortex-trained clocks (7.74, 95% CI [1.89, 31.69]) showed substantially higher prediction error. It is notable that confidence intervals were relatively wide for all training tissues.

Leave-one-cohort-out analyses indicated that the tissue-training pattern was robust across cohort exclusions, with tissue-specific predicted geometric mean $MAE_w$ values remaining relatively stable across re-fitted models, particularly for cord-trained clocks (range: 0.04–0.11) and multi-tissue-trained clocks (range: 1.51–2.02)

*Tissue of test sample*

Epigenetic age prediction performance differed significantly across test tissues (QM(5) = 74.10, $p_{FDR}$ < .001), with the lowest predicted geometric mean $MAE_w$ observed in blood (1.51, 95% CI [0.56, 4.10]) and saliva samples (1.46, 95% CI [0.53, 4.01]), intermediate error in buccal samples (4.33, 95% CI [1.26, 14.92]), and substantially higher error in cord (13.27, 95% CI [4.41, 39.89]) and placenta samples (24.73, 95% CI [5.43, 112.62]). The markedly elevated prediction error in cord and placenta samples likely reflects that non-gestational clocks were also applied to the birth timepoints with cord and placenta tissues. It is also of note that only one sample had placenta data.

Leave-one-cohort-out analyses indicated that the tissue-specific performance pattern was generally robust across cohort exclusions, with relatively stable estimates for blood, saliva, and cord samples, although buccal estimates showed greater variability (predicted geometric mean $MAE_w$ range: 2.46–13.06).

*Pearson correlations*

*Tissue overlap*

Epigenetic clocks applied to matching tissue types showed significantly stronger correlations with chronological age than clocks applied to non-matching tissues ($b_z$ = 0.11 Fisher-z units, 95% CI [0.093, 0.118], $p_{FDR}$ < .001). Predicted pooled correlations increased from $r$ = .31 for non-matching tissues to $r$ = .42 for matching tissues, indicating a modest but reliable improvement in performance when training and test tissues aligned.

Tissue overlap explained a significant proportion of between-study heterogeneity (QM(1) = 264.61, $p_{FDR}$ < .001), although substantial residual heterogeneity remained (QE(733) = 22502.57, $p$ < .001; $I^2$ = 98.35%). Variance decomposition indicated that heterogeneity was primarily attributable to differences between clock models (51.56% of observed variance), followed by cohorts (41.38%) and developmental time points within cohorts (5.40%).

Leave-one-cohort-out analyses demonstrated that the tissue-overlap effect was robust and consistently positive across all re-fitted models ($b_z$ range = 0.08–0.13 Fisher-z units; SD = 0.01).

*Tissue of the training sample*

Training tissue significantly moderated epigenetic clock performance (QM(5) = 129.50, $p_{FDR}$ < .001), with cord-blood-trained clocks showing by far the strongest correspondence with chronological age ($r$ = .75, 95% CI [.66, .82]). In contrast, clocks trained on blood ($r$ = .24, 95% CI [.10, .37]), cortex ($r$ = .24, 95% CI [.04, .42]), buccal tissue ($r$ = .25, 95% CI [.003, 0.472]) and multi-tissue datasets ($r$ = .32, 95% CI [.15, .47]) showed substantially weaker correlations overall. Buccal-trained clocks showed comparatively wide confidence intervals.

Despite the significant effect of training tissue, substantial heterogeneity remained across estimates (QE(730) = 19856.79, $p$ < .001; $I^2$ = 97.27%). Variance decomposition indicated that most variability was attributable to cohort differences (71.61% of observed variance), followed by clock-model differences (16.12%) and developmental time points within cohorts (9.54%).

Leave-one-cohort-out analyses demonstrated that tissue-specific findings were generally robust across cohort exclusions. Cord-trained clocks consistently showed the strongest performance (pooled $r$ range = .63–.78), although estimates were somewhat attenuated when GenR was excluded. Performance estimates for blood-, cortex-, and multi-tissue-trained clocks remained highly stable across cohort exclusions (all SDs ≤ .023), indicating that no single cohort disproportionately influenced the observed tissue-training patterns

*Tissue of test sample*

Test tissue significantly moderated epigenetic clock performance (QM(5) = 21.54, $p_{FDR}$ = .001). Clocks applied to buccal samples showed the strongest overall correspondence with chronological age ($r$ = .47, 95% CI [.25, .64]), followed by blood ($r$ = .38, 95% CI [.20, .53]), cord blood ($r$ = .33, 95% CI [.13, .50]) and saliva ($r$ = .30, 95% CI [.12, .47]). In contrast, clocks applied to placental tissue showed comparatively weak and non-significant associations with chronological age ($r$ = .0.20, 95% CI [-0,10, .46], $p_{uncorrected}$ = .190).

Despite significant tissue-specific differences, substantial heterogeneity remained across estimates (QE(730) = 22577.71, $p$ < .001; $I^2$ = 98.63%). Variance decomposition indicated that heterogeneity was primarily attributable to clock-model differences (51.31% of observed variance), followed by cohorts (43.15%), whereas developmental time points within cohorts accounted for comparatively little variability (4.16%).

Leave-one-cohort-out analyses demonstrated generally robust tissue-specific findings across cohort exclusions. Estimates for blood, cord and saliva samples remained highly stable (all SDs ≤ .032). Buccal estimates showed greater variability across cohort exclusions (*r* range = .16-.52), largely driven by the exclusion of UCI Echo which reduced the age correlation, indicating some sensitivity of buccal-tissue estimates to individual cohorts. Placenta data was only available in one cohort.

#### SM 2.5 Performance of brain age models across development

*Unweighted MAE*

Across all brain age models, cohorts, and developmental periods, the pooled MAE was 4.46 years (95% CI [2.48, 6.43]), indicating moderate overall prediction error across childhood and adolescence. Heterogeneity was extremely high ($I^2$ = 99.90%) and was primarily attributable to differences between brain age models (85.92%), with comparatively smaller contributions from cohorts (10.57%) and developmental time points within cohorts (3.41%). Leave-one-cohort-out analyses showed stable pooled estimates (MAE range = 4.34–4.71 years), suggesting that no individual cohort disproportionately influenced the overall findings.

Brain age models differed substantially in prediction accuracy (QM(7) = 31070.95, $p_{\text{uncorrected}}$ < .001), with the lowest MAEs observed for Pyment (2.20 years), DevBrainAge (2.61 years), and Centile2 (2.61 years), whereas ENIGMA (8.00 years) and PyBrainAge (8.23 years) showed the largest prediction errors. Residual heterogeneity remained very high after accounting for model differences ($I^2$ = 99.30%), with most remaining variability attributable to cohorts (75.09%) rather than developmental time points (24.21%). Modelwise leave-one-cohort-out analyses demonstrated stable performance estimates across cohorts for most models, indicating that the relative ranking of brain age models was robust to the exclusion of individual cohorts. The largest differences were observed for DBN (MAE range = 2.90 - 4.89, exclusion of Oregon ADHD-1000 or GUSTO increased MAE) and ENIGMA (MAE range = 7.50 - 9.66, exclusion of GenR increases MAE).

*Weighted MAE in a restricted subsample*

We reran the main brain age performance ($MAE_w$) meta-analysis after restricting it to effect sizes derived from samples/timepoints where the age ranges of the test data and training data overlapped. This subset included 60.9% of the data of the main $MAE_w$ and retained all brain age models (except DunedinPACNI which was not included in the $MAE_w$ analyses throughout this study).

Even though we found that age range overlap between training and test data was associated with a lower $MAE_w$ (see SM 2.7), the pooled $MAE_w$ in this subsample was slightly higher (2.03, 95% CI [0.37, 3.69]) than in the full sample analyses (1.72). Even in this 'cleaner' subsample, heterogeneity was high ($I^2$ = 100, $p$ < .001).

*Pearson correlations*

The overall pooled Pearson correlation between predicted brain age and chronological age across all brain age models, cohorts and developmental periods was small-to-moderate in magnitude ($r$ = .20, 95% CI [.07, .32]), indicating modest overall correspondence between brain age estimates and chronological age across development. Substantial heterogeneity was observed across estimates ($I^2$ = 93.84%, Q(214) = 3659.36, $p$ < .001). Variance decomposition indicated that heterogeneity was primarily attributable to cohort differences (48.11%) and clock-model differences (41.70%), whereas developmental time point within cohorts accounted for comparatively little variability (4.04%). Leave-one-cohort-out analyses demonstrated high robustness of the pooled findings, with pooled correlations remaining highly stable across cohort exclusions (pooled $r$ range = .17–.21).

Modelwise analyses revealed substantial differences in chronological-age correspondence across brain models (QM(8) = 1481.05, $p$ < .001). The strongest pooled correlation was observed for Pyment ($r$ = .33, 95% CI [.25, .40]) and the weakest for ENIGMA ($r$ = .10, 95% CI [.01, .18]). DunedinPACNI was the only clock showing a significant negative pooled correlation with chronological age ($r$ = -.10, 95% CI [-.19, -.02]). Leave-one-cohort-out analyses indicated that model-specific findings were highly stable across cohort exclusions (SDs < 0.025), including for DunedinPACNI, whose pooled correlations remained consistently negative across all leave-one-cohort-out analyses.

### SM 2.6 Brain age performance moderators: Developmental period

*Pearson correlations*

In keeping with the $MAE_w$ results, the overall association between predicted brain age and chronological age did not significantly change across childhood and adolescence ($b_z$ = 0.005, $p_{FDR}$ = .241), indicating relatively stable pooled correlations across development. Nevertheless, substantial residual heterogeneity remained (QE(213) = 3612.24, $p$ < .001; $I^2$ = 93.45%). Variability was distributed relatively evenly between cohort-level differences (45.06% of heterogeneity) and clock-model differences (44.07%), whereas developmental time point within cohorts explained comparatively little heterogeneity (4.32%). Leave-one-cohort-out analyses demonstrated robustness of the (lack of a) global developmental effect, with slope estimates ($b$) ranging from -0.002 to 0.007).

Despite the absence of a significant overall developmental trend, modelwise analyses revealed differences in developmental trajectories across brain age models (QM(16) = 1618.41, $p_{uncorrected}$ < .001). Allowing developmental slopes to vary across models substantially reduced residual heterogeneity (QE(199) = 1751.24, $p$ < .001) and lowered total heterogeneity from 93.45% to 88.00%.

Some brain age models showed increasingly stronger chronological-age correspondence in older developmental samples, including DunedinPACNI ($b_z$ = 0.02, 95% CI [0.01, 0.03], $p_{FDR}$ < .001) and ENIGMA ($b_z$ = 0.01, 95% CI [0.00, 0.02], $p_{FDR}$ = .058). In contrast, DevBrainAge ($b_z$ = -0.017, 95% CI [-0.03, -0.01], $p_{FDR}$ = .001) and PyBrainAge ($b_z$ = -0.01, 95% CI [-0.02, -

0.00], $p_{FDR}$ = .053) showed weaker correlations with chronological age in older samples. Other clocks, including Centile2, DBN, Kaufmann and Pyment, showed comparatively stable developmental associations with chronological age.

Leave-one-cohort-out analyses demonstrated robustness of the model-specific developmental effects, indicating minimal influence of individual cohorts on the observed developmental trajectories (SDs < 0.01).

DunedinPACNI differed markedly from the other brain models by showing an overall negative pooled correlation with chronological age ($r$ = -.10, 95% CI [-.19, -.02]). However, DunedinPACNI also showed the strongest positive developmental age slope ($b_z$ = 0.02, $p_{FDR}$ < .001, indicating that its association with chronological age increased substantially across development.

**SM 2.7 Brain age performance moderators: Age range overlap between brain age training and testing data**

*Weighted MAE*

Age overlap was defined as the percentage of the test sample age range that is covered by the training age range (overlap / study age range).

A greater proportional overlap between the cohort age range and the brain model training age range was associated with significantly lower $MAE_w$, with each 1% increase in overlap linked to an average ~0.0037 reduction in $MAE_w$ ($b$ = -0.0033, 95% CI [-0.0037, -0.0030], $p_{FDR}$ < .001). Predicted $MAE_w$ decreased from approximately 2.01 at 0% overlap to 1.67 at 100% overlap, indicating progressively better apparent model performance with increasing training-age overlap. Proportional age overlap explained a substantial proportion of between-study heterogeneity (QM(1) = 365.30, $p_{FDR}$ < .001), and leave-one-cohort-out analyses showed that the effect was highly robust, remaining consistently negative across all re-fitted models ($b$ range: -0.0047 to -0.0030; SD = 0.0004).

*Pearson correlations*

Greater proportional overlap between the training-age range and the test-sample age range was associated with significantly stronger correlations between predicted brain age and chronological age ($b_z$ = 0.0014 Fisher-z units per 1% overlap increase, 95% CI [0.0010, 0.0019], $p_{FDR}$ < .001). Predicted pooled correlations increased from $r$ = .10 in samples with no overlap to $r$ = .24 in samples with complete (100%) overlap between training and test age ranges.

Proportional age overlap explained a substantial proportion of between-study heterogeneity (QM(1) = 44.11, $p_{FDR}$ < .001), although considerable residual heterogeneity remained across cohorts, timepoints and models (QE(213) = 2422.21, $p$ < .001; $I^2$ = 91.65%). Variance decomposition indicated that heterogeneity was primarily attributable to cohorts (58.45%), followed by clock models (27.92%) and developmental time points within cohorts (5.27%).

Leave-one-cohort-out analyses demonstrated that the estimated effect remained consistently positive across all re-fitted models ($b_z$ range = 0.0005–0.0017 Fisher-z units per 1% overlap increase; SD = 0.00028) although the magnitude of the association was attenuated when ALSPAC was excluded ($b$ = 0.0005), suggesting some influence of this cohort on the overall effect estimate.

**SM 2.8 Brain age performance moderators: Processing level (minimal vs extracted)**

*Weighted MAE*

Although brain age models based on extracted imaging-derived features showed numerically higher $MAEs_w$ than models based on minimal processing (predicted $MAE_w$: 1.83 vs 1.43), the estimated effect was small and non-significant, $b_{log}$ = 0.41, 95% CI [-0.33, 1.14], $p_{FDR}$ = .369), indicating no clear evidence that “minimal” versus “extracted” processing pipelines were associated with differences in $MAE_w$. The processing moderator did not explain a significant proportion of between-study heterogeneity (QM(1) = 1.18, $p_{FDR}$ = .369), and substantial residual heterogeneity remained across cohorts, timepoints and models (QE(182) = 19388.43, $p$ < .001; $I^2$ = 100.00%). Variance decomposition indicated that heterogeneity was primarily attributable to cohort differences (94.50%), whereas clock-model differences (5.27%) and developmental time points within cohorts (0.19%) accounted for comparatively little variability.

Leave-one-cohort-out analyses suggested that the processing effect was less stable than the age-overlap moderators, with the estimated effect varying substantially across cohort exclusions and even reversing direction when Oregon ADHD-1000 was removed ($b_{log}$ range: -0.54 to 0.45, SD = 0.28).

*Pearson correlations*

Processing level was not significantly associated with brain-age performance measured using Pearson’s correlation coefficient with chronological age ($b_z$ = -0.13 Fisher-z units, 95% CI [-0.36, 0.10], $p_{FDR}$ = .258), indicating that extracted processing pipelines did not show reliably different chronological-age correlations compared with minimally processed data. Predicted pooled correlations were $r$ = .29 for minimally processed data and $r$ = .16 for extracted data.

The processing moderator did not explain a significant proportion of between-study heterogeneity (QM(1) = 1.28, $p_{FDR}$ = .258), and substantial residual heterogeneity remained across cohorts, timepoints and models (QE(213) = 3272.71, $p$ < .001; $I^2$ = 93.65%). Variance decomposition indicated that heterogeneity was primarily attributable to cohort differences (48.88%) and clock-model differences (40.67%), whereas developmental time points within cohorts accounted for comparatively little variability (4.10%).

Leave-one-cohort-out analyses demonstrated robustness of the processing effect estimates, with the moderator effect remaining consistently negative and highly stable across all re-fitted models ($b_z$ range = -0.141 to -0.121 Fisher-z units; SD = 0.005). However, confidence intervals consistently crossed zero, supporting the conclusion.

#### SM 2.9 Within-modality associations

Within-modality correlations (without covariate adjustments) were mostly positive across both brain and epigenetic aging modalities (SM Figures S2 and S3), indicating moderate shared variance between models within each modality. Global pooled correlations were:

- Brain predicted age: *r* = .29 (95% CI: .21–.37)
    - Model-combination correlations ranged from *r* = −.165 to *r* = .48
    - Largest positive correlations:
        - Kaufmann–PyBrainAge (*r* = .477)
        - DBN–Pyment (*r* = .468)
        - Kaufmann–Pyment (*r* = .450)
        - ENIGMA–Kaufmann (*r* = .435)
        - Centile2–Kaufmann (*r* = .433).
    - Largest negative correlations:
        - DevBrainAge–DunedinPACNI (*r* = −.165)
        - DunedinPACNI–PyBrainAge (*r* = −.069)
        - Centile2–DunedinPACNI (*r* = −.048)
- Brain PAR: *r* = .25 (95% CI: .19–.32)
    - Model-combination correlations ranged from *r* = −.098 to *r* = .443
    - Largest positive correlations:
        - Kaufmann–PyBrainAge (*r* = .443)
        - DBN–Pyment (*r* = .414)
        - ENIGMA–Kaufmann (*r* = .395)
        - DBN–PyBrainAge (*r* = .389)
        - DBN–DevBrainAge (*r* = .380).
    - Largest negative correlations (included DunedinPACNI):
        - DevBrainAge–DunedinPACNI (*r* = −.098)
        - DunedinPACNI–PyBrainAge (*r* = −.046)
- Epigenetic predicted age: *r* = .32 (95% CI: .24–.39)
    - Model-combination correlations showed a much wide*r* range, from *r* = −.517 to *r* = .936.
    - Largest positive correlations:
        - Bohlin–EPIC (*r* = .936)
        - Bohlin–Knight (*r* = .764)
        - EPIC–Knight (*r* = .739)
        - cAge–ZhangEN (*r* = .706)
        - DNAmTL–PCGrimAge (*r* = .661).
    - Largest negative correlations:
        - AdaptAge–DamAge (*r* = −.527)
        - AdaptAge–DNAmTL (*r* = −.167)
        - AdaptAge–cAge (*r* = −.103)
        - AdaptAge–Hannum (*r* = −.088)
- Epigenetic PAR: *r* = .22 (95% CI: .18–.26)
    - Model-combination correlations ranged from *r* = −.574 to *r* = .813

- Largest positive correlations:
  - Bohlin–EPIC ($r$ = .813)
  - cAge–ZhangEN ($r$ = .631)
  - DNAmTL–PCGrimAge ($r$ = .621)
  - cAge–DamAge ($r$ = .610)
  - DNAmTL–Hannum ($r$ = .591)
- Largest negative correlations - these all included AdaptAge, which aims to estimate protective causal aging influences:
  - AdaptAge–DamAge ($r$ = −.517)
  - AdaptAge–DNAmTL ($r$ = −.155)
  - AdaptAge–cAge ($r$ = −.091)
  - AdaptAge–ZhangEN ($r$ = −.062)

Overall, brain model combinations showed moderate and relatively constrained within-modality correlations (with negative correlations involving DunedinPACNI), whereas epigenetic model combinations showed much broader heterogeneity, including both strong positive correlations among closely related clocks and negative correlations involving AdaptAge.

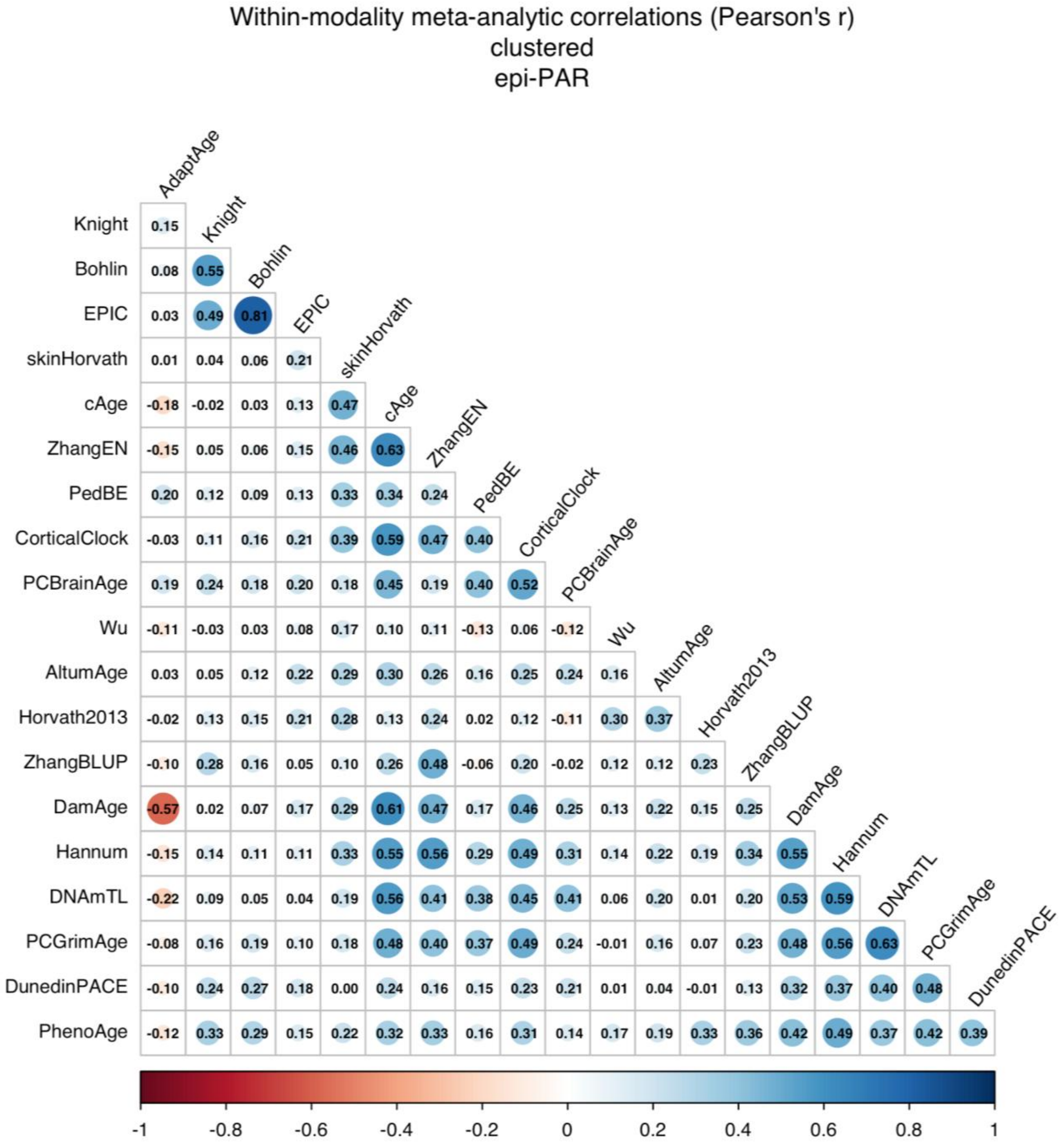


**SM Figure S2. Within-modality correlations of epigenetic PARs.**

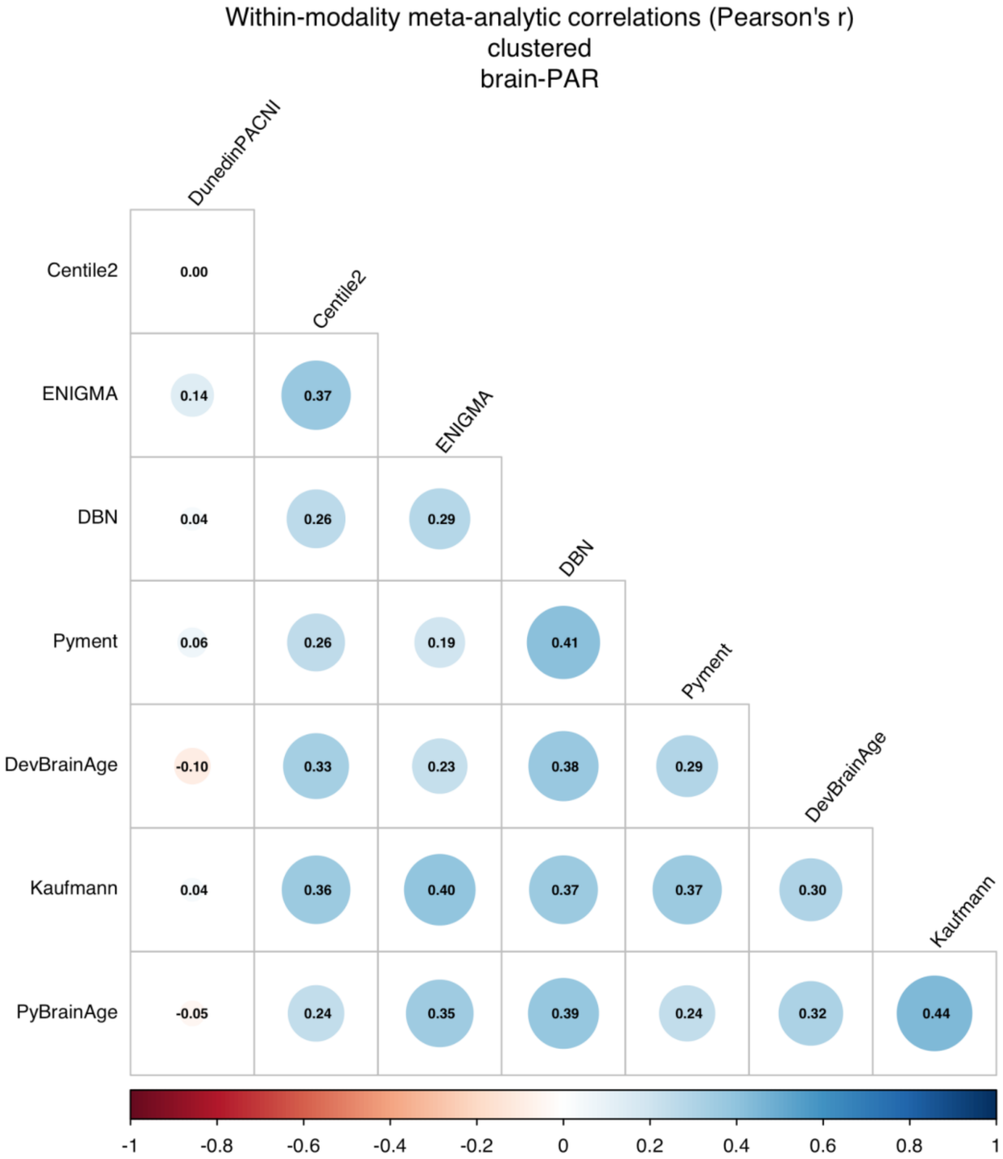


**SM Figure S3. Within-modality correlations of brain age PARs.**

### SM 2.10 The influence of epigenetic clock generation on brain PAR - epigenetic PAR associations

*Epigenetic clock generation (with 2 levels: 1st vs next-generation)*
Brain PAR–epigenetic PAR associations did not significantly differ between first-generation epigenetic clocks and later-generation clocks overall (QM(1) = 0.15, $p_{FDR}$ = .700). The estimated standardized association beta for first-generation clocks was 0.016 (95% CI [0.006, 0.026]), while later-generation clocks showed only a small, non-significant increase in association strength relative to first-generation clocks (β difference = 0.0014, 95% CI [-0.0055, 0.0083]). Leave-one-cohort-out analyses indicated that the null generation effect was robust across cohort exclusions, with estimated NextGen-versus-Gen1 differences

remaining consistently close to zero across all re-fitted models (β difference range: -0.00002–0.00334; SD = 0.0009).

*Epigenetic clock generation (with 4 levels: 1st, 2nd, 3rd, 4th)*
Brain PAR–epigenetic PAR associations differed across epigenetic clock generations (QM(4) = 20.11, $p_{FDR}$ = .002), with the strongest associations observed for DunedinPACE (the only third generation clock). Estimated standardized association betas were 0.016 (95% CI [0.006, 0.026]) for first-generation clocks, 0.018 (95% CI [0.006, 0.031]) for second-generation clocks, 0.033 (95% CI [0.017, 0.049]) for DunedinPACE (i.e., the only third-generation clock), and 0.010 (95% CI [-0.003, 0.022]) for fourth-generation clocks, with only the fourth-generation estimate not significantly different from zero (uncorrected). Leave-one-cohort-out analyses indicated that the generation-specific association pattern was robust across cohort exclusions, with relatively small variability in estimated effects across re-fitted models (SD range: 0.0016–0.0030).

**SM 2.11 The influence of tissue on brain PAR - epigenetic PAR associations**

*Saliva in either test or training data*
Saliva involvement in either the test sample or the epigenetic clock training data did not significantly moderate brain PAR–epigenetic PAR associations (QM(1) = 1.19, *b* = 0.004, 95% CI [-0.004,0.012], $p_{FDR}$ = .477). Predicted standardized association betas were very similar for no saliva involvement (β = 0.015, 95% CI [0.005, 0.025]) versus when saliva was involved (β = 0.019, 95% CI [0.009, 0.030]). Leave-one-cohort-out analyses supported this null finding: the saliva-related difference remained small across all cohort exclusions (*b* range: 0.0006–0.0072; SD = 0.0017), with all confidence intervals crossing zero. Results were consistent when saliva involvement was defined as saliva tissue in test *and* training data (QM(1) = 0.45, *b* = 0.005, 95% CI [-0.009,0.019], $p_{uncorrected}$ = .500). Notably, for saliva to be considered involved in training, the clock did not have to be exclusively trained on saliva (e.g., the multi-tissue clock Horvath2013 would be classified as containing saliva in the training data).

*Buccal in either test or training data*
Buccal involvement in either the test sample or the epigenetic clock training data did not significantly moderate brain PAR–epigenetic PAR associations (QM(1) = 0.87, *b* = 0.004, 95% CI [-0.004,0.011], $p_{FDR}$ = .477). Predicted standardized association betas were highly similar for non-buccal and buccal-involved comparisons: β = 0.016 (95% CI [0.006, 0.026]) for no buccal involvement versus β = 0.019 (95% CI [0.008, 0.030]) when buccal tissue was involved. Leave-one-cohort-out analyses supported this null finding, with the estimated buccal-related difference remaining consistently small across all cohort exclusions (*b* range: 0.001–0.005; SD = 0.001), and all confidence intervals overlapping zero. Results were consistent when buccal involvement was defined as saliva tissue in test *and* training data (QM(1) = 1.47, *b* = 0.011, 95% CI [-0.007,0.029], $p_{uncorrected}$ = .225). However, these analyses relied on one cohort (GUSTO) only. Notably, for buccal to be considered involved in training, the clock did not have to be exclusively trained on buccal (e.g., Horvath2013).

*Cortex tissue in training data*
Cortical tissue involvement in epigenetic clock training data showed a small, non-significant tendency toward stronger brain PAR–epigenetic PAR associations (QM = 3.68, $p_{FDR}$ = .128). Predicted standardized association betas were β = 0.015 (95% CI [0.005, 0.025]) for non-cortical comparisons and β = 0.022 (95% CI [0.011, 0.033]) when cortical tissue was involved, corresponding to an estimated increase of 0.007 in association strength that did not reach statistical significance (95% CI [-0.0002,0.0148], $p_{FDR}$ = .128). Leave-one-cohort-out analyses suggested that the positive cortical-tissue effect was relatively stable across cohort exclusions (difference range: 0.004–0.009; SD = 0.001); 7/12 confidence intervals did not include 0.

**SM 2.12 Sensitivity analyses of brain PAR–epigenetic PAR associations under alternative covariate adjustment**

The primary model to assess for brain PAR - epigenetic PAR associations at the cohort level corrected for age at MRI, age at DNA methylation, sex and batch. To evaluate the robustness of the primary findings, we repeated the analyses using two alternative covariate adjustment strategies: (1) no covariate adjustment and (2) covariates from the primary model and additional adjustment for estimated cell type proportions. Summary results are presented in SM Table 2.12.1.

Across both sensitivity analyses, the pooled associations between brain PAR and epigenetic PAR were highly consistent with the primary analyses. The estimated effect size remained virtually unchanged (β = 0.017 in both models), although statistical significance was attenuated after adjustment for cell type proportions (SM Table 2.12.1). Residual heterogeneity also remained modest and comparable to that observed in the primary analyses.

The pattern of results across individual brain age–epigenetic age model combinations was similarly stable. Approximately three-quarters of pooled associations remained positive, more than 80% of model combinations retained the same direction of effect as in the primary analyses, and effect estimates were strongly correlated with those from the primary analyses (*r* = 0.71 without covariates; *r* = 0.85 with additional cell type adjustment). Together, these findings indicate that the primary results were relatively robust to alternative covariate adjustment strategies.

**SM Table S2.12.1. Brain PAR–epigenetic PAR associations under alternative covariate adjustment strategies**

| | Model | Primary | No covariates | Cell Types |
|---|---|---|---|---|
| Global | Pooled β (95% CI) | 0.017 (0.01, 0.03) | 0.017 (0.01, 0.03) | 0.017 (-.002, 0.036) |
| | *p* | .001 | .001 | .083 |
| | $I^2$ | 14.64% | 18.10% | 19.10% |
| | Q *p* | 0.578 | < .001 | 0.975 |
| Model combi | QM *p* | < .001 | < .001 | < .001 |
| | Positive associations | 77.90% | 77.21% | 75.74% |
| | Negative associations | 22.10% | 22.79% | 24.26% |
| | Same sign as primary | - | 83.09% | 84.56% |
| | Correlation with primary βs | - | 0.71 | 0.85 |

Note: Q *p* = statistical significance of the Q-test for heterogeneity, QM *p* = statistical significance of Q-test for the moderating effect of model combinations.

**SM 2.13 Brain PAR - epigenetic PAR associations in a restricted subsample**

Restricting the analyses to model combinations in which the test sample age range was fully contained within the training age range and epigenetic clocks were applied to their intended tissue reduced the dataset to 385 associations (13.7% of all associations), spanning 47 model combinations, 7 brain age models, and 7 epigenetic age models.

In this restricted dataset, the association between brain PAR and epigenetic PAR remained small and non-significant (β = 0.012, 95% CI [-0.007, 0.031], *p* = .218), closely matching the effect observed in the main analyses (β = 0.017). The lack of statistical significance likely reflects the substantially reduced sample size. Mirroring the analyses of the full sample, heterogeneity was non-significant, Q(384) = 382.7140, *p* = .509, $I^2$ = 33.7.

Similarly, the moderating effect of sample mean age on brain–PAR - epigenetic PAR associations remained small and non-significant (*b* = -0.002, 95% CI [-0.007, 0.004], *p* = .551), suggesting that epigenetic-brain age associations did not depend on the developmental stage of the sample. Although the direction of the effect differed from that observed in the main analyses (β = 0.001), the estimated effects were close to zero in both analyses.

**References for all included biological age models**

| First author | Name (current study) | Full reference |
|---|---|---|
| **Epigenetic clocks** | | |
| Belsky 2022 | DunedinPACE | Belsky, D. W. *et al.* DunedinPACE, a DNA methylation biomarker of the pace of aging. *eLife* **11**, e73420 (2022). |
| Bernabeu 2023 | cAge | Bernabeu, E. *et al.* Refining epigenetic prediction of chronological and biological age. *Genome Med.* **15**, 12 (2023). |
| Bohlin 2016 | Bohlin | Bohlin, J. *et al.* Prediction of gestational age based on genome-wide differentially methylated regions. *Genome Biol.* **17**, 207 (2016). |
| de Lima Camillo 2022 | AltumAge | de Lima Camillo, L. P., Lapierre, L. R. & Singh, R. A pan-tissue DNA-methylation epigenetic clock based on deep learning. *npj Aging* **8**, 4 (2022). |
| Haftorn 2021 | EPIC | Haftorn, K. L. *et al.* An EPIC predictor of gestational age and its application to newborns conceived by assisted reproductive technologies. *Clin. Epigenetics* **13**, 82 (2021). |
| Hannum 2013 | Hannum | Hannum, G. *et al.* Genome-wide methylation profiles reveal quantitative views of human aging rates. *Mol. Cell* **49**, 359–367 (2013). |
| Higgins-Chen 2022 | PCGrimAge | Higgins-Chen, A. T. *et al.* A computational solution for bolstering reliability of epigenetic clocks: implications for clinical trials and longitudinal tracking. *Nat. Aging* **2**, 644–661 (2022). |
| Horvath 2013 | Horvath2013 | Horvath, S. DNA methylation age of human tissues and cell types. *Genome Biol.* **14**, 3156 (2013). |

| | | |
|---|---|---|
| Horvath 2018 | skinHorvath | Horvath, S. *et al.* Epigenetic clock for skin and blood cells applied to Hutchinson–Gilford Progeria syndrome and *ex vivo* studies. *Aging (Albany NY)* **10**, 1758–1775 (2018). |
| Knight 2016 | Knight | Knight, A. K. *et al.* An epigenetic clock for gestational age at birth based on blood methylation data. *Genome Biol.* **17**, 206 (2016). |
| Levine 2018 | PhenoAge | Levine, M. E. *et al.* An epigenetic biomarker of aging for lifespan and healthspan. *Aging (Albany NY)* **10**, 573–591 (2018). |
| Lu 2019 | DNAmTL | Lu, A. T. *et al.* DNA methylation-based estimator of telomere length. *Aging (Albany NY)* **11**, 5895–5923 (2019). |
| McEwen 2020 | PedBE | McEwen, L. M. *et al.* The PedBE clock accurately estimates DNA methylation age in pediatric buccal cells. *Proc. Natl Acad. Sci. USA* **117**, 23329–23335 (2020). |
| Shireby 2020 | CorticalClock | Shireby, G. L. *et al.* Recalibrating the epigenetic clock: implications for assessing biological age in the human cortex. *Brain* **143**, 3763–3775 (2020). |
| Thrush 2022 | PCBrainAge | Thrush, K. L. *et al.* Aging the brain: multi-region methylation principal component based clock in the context of Alzheimer's disease. *Aging (Albany NY)* **14**, 5641–5668 (2022). |
| Wu 2019 | Wu | Wu, X. *et al.* DNA methylation profile is a quantitative measure of biological aging in children. *Aging (Albany NY)* **11**, 10031–10051 (2019). |
| Ying 2024 | AdaptAge | Ying, K. *et al.* Causality-enriched epigenetic age uncouples damage and adaptation. *Nat. Aging* **4**, 231–246 (2024). |
| Ying 2024 | DamAge | Ying, K. *et al.* Causality-enriched epigenetic age uncouples damage and adaptation. *Nat. Aging* **4**, 231–246 (2024). |
| Zhang 2019 | ZhangEN | Zhang, Q. *et al.* Improved precision of epigenetic clock estimates across tissues and its implication for biological ageing. *Genome Med.* **11**, 54 (2019). |
| Zhang 2019 | ZhangBLUB | Zhang, Q. *et al.* Improved precision of epigenetic clock estimates across tissues and its implication for biological ageing. *Genome Med.* **11**, 54 (2019). |
| **Brain age models** | | |
| Bashyam 2020 | DeepBrainNet | Bashyam, V. M. *et al.* MRI signatures of brain age and disease over the lifespan based on a deep brain network and 14,468 individuals worldwide. *Brain* **143**, 2312–2324 (2020). |
| Cole 2023 | PyBrainAge | Cole, J. *PyBrainAge*. GitHub (2023). |
| Drobinin 2022 | DevBrainAge | Drobinin, V. *et al.* The developmental brain age is associated with adversity, depression, and functional outcomes among adolescents. *Biol. Psychiatry Cogn. Neurosci. Neuroimaging* **7**, 406–414 (2022). |
| Han 2021 | ENIGMA | Han, L. K. M. *et al.* Brain aging in major depressive disorder: results from the ENIGMA major depressive disorder working group. *Mol. Psychiatry* **26**, 5124–5139 (2021). |
| Kaufmann 2019 | Kaufmann | Kaufmann, T. *et al.* Common brain disorders are associated with heritable patterns of apparent aging of the brain. *Nat. Neurosci.* **22**, 1617–1623 (2019). |

| | | |
|---|---|---|
| Leonardsen 2022 | Pyment | Leonardsen, E. H. *et al.* Deep neural networks learn general and clinically relevant representations of the ageing brain. *Neuroimage* **256**, 119210 (2022). |
| Whitman 2025 | DunedinPACNI | Whitman, E. T. *et al.* DunedinPACNI estimates the longitudinal pace of aging from a single brain image to track health and disease. *Nat. Aging* **5**, 1619–1636 (2025). |
| Yu 2024 | Centile2 | Yu, Y. *et al.* Brain-age prediction: systematic evaluation of site effects, and sample age range and size. *Hum. Brain Mapp.* **45**, e26768 (2024). |
